# Sex without crossovers mimics clonal reproduction in the holocentric plant *Rhynchospora tenuis*

**DOI:** 10.64898/2026.01.17.700054

**Authors:** Meng Zhang, Marco Castellani, Stefan Steckenborn, Maciej Majka, Georgios Tsipas, Thiago Nascimento, Ulla Neumann, Gokilavani Thangavel, Laura A. Robledillo, Thomas Lux, Lorraine Deberón, Ursula Pfordt, José A. Campoy, Athul Vijayan, Ton Timmers, Nafiseh Sargheini, Magdalena Marek, Hequan Sun, Paulo G. Hofstatter, Bruno Huettel, Steven Dreissig, Klaus F. X. Mayer, Duarte D. Figueiredo, André L. L. Vanzela, Korbinian Schneeberger, André Marques

## Abstract

Meiotic recombination ensures accurate chromosome segregation and promotes genetic diversity by generating crossovers between homologous chromosomes^1^. While essential in most sexually reproducing organisms, recombination is variably regulated and can be absent in some lineages, a condition known as achiasmy^2^. However, obligate achiasmy in both sexes of a sexual species has not been previously documented. Here, we investigate the beak-sedge *Rhynchospora tenuis*, a holocentric plant with the lowest known chromosome number among flowering plants (*n* = 2) and inverted meiosis^3^. Using chromosome-scale genome assemblies from nine accessions, molecular cytogenetics, immunocytochemistry, high-throughput single-gamete sequencing and whole-genome sequencing of controlled crosses, we show that *R. tenuis* undergoes obligate, genome-wide achiasmy in both male and female meiosis. Despite normal early meiotic axis formation, synapsis fails, crossovers are not detected cytologically or genetically, and univalents persist at metaphase I. Extensive haplotype-specific accumulation of transposable elements (TEs) generates segregation distortion (e.g. meiotic drive), favouring the transmission of larger, TE-rich chromosomes. Remarkably, sexual reproduction is retained with fertilisation producing viable seeds only when translocation-compatible gametes meet, indicating strong post-meiotic selection that eliminates incompatible homozygous combinations. As a result, all surviving offspring are genetically identical to the maternal genotype, effectively restoring heterozygosity each generation and mimicking clonal reproduction. We propose that the combined effects of recombination loss, low chromosome number, holocentricity, inverted meiosis, and selective transmission of longer chromosomes enable faithful segregation and clonal-like inheritance despite sexual reproduction. These findings challenge the boundary between sex and clonality, revealing a unique evolutionary strategy linking genome architecture, recombination loss, and transmission bias.

## Introduction

Although meiotic recombination is critical for most sexually reproducing species, its frequency varies widely, often differing between sexes (heterochiasmy)^4,5^, and can be entirely suppressed in specific chromosomes or lineages, known as achiasmy^2^. In extreme cases, such as male *Drosophila melanogaster*, meiotic recombination and chiasmata are completely absent, requiring alternative mechanisms like chromatin threads or meiotic drive to ensure correct chromosome segregation^6–10^. The absence of recombination is expected to have profound consequences, including impaired chromosome segregation, reduced genetic diversity, and accelerated mutation accumulation through the so-called ‘Meselson effect’^11–14^. However, obligate, genome-wide bisexual achiasmy of a sexually reproducing organism has never been described.

The beak-sedge *Rhynchospora tenuis* Link (*n* = 2) presents a compelling model to investigate these questions. It harbours only two pairs of holocentric chromosomes, the lowest known number in plants, and undergoes inverted meiosis, with sister chromatids separating at meiosis I, followed by the segregation of homologous chromosomes at meiosis II^3,15^. Its holocentric chromosomes, in which kinetochore activity is distributed along their entire length, are known to tolerate structural rearrangements such as chromosome end-to-end translocations (hereafter called ‘fusions’)^16^, which may have facilitated the evolution of its extremely reduced chromosome number^17^. Previous cytological studies revealed that male meiosis in *R. tenuis* proceeds without homologous chromosome pairing and chiasmata formation, displaying four univalents at diakinesis, raising the possibility of achiasmy despite apparent meiotic DNA double-strand breaks (DSB) formation^3^. However, it remains unclear whether and how meiotic recombination is completely abolished, particularly during female meiosis, and whether genome transmission remains stable in the absence of crossovers (COs).

Here, we investigate the recombination landscape, meiotic progression, and inheritance patterns of *R. tenuis* using an integrative approach combining chromosome-scale pangenomics, molecular cytogenetics and immunocytochemistry, single-gamete sequencing of pollen nuclei, plus whole-genome sequencing of offspring from controlled crosses. We demonstrate that *R. tenuis* exhibits obligate bisexual achiasmy, with complete absence of recombination, synapsis defects, and striking segregation distortion favouring larger chromosomes, enriched with TEs. Remarkably, this results in progeny that mirrors the mother plant genotype despite sexual reproduction. Our findings uncover a reproductive system in which meiotic drive and post-meiotic selection effectively replace the need for recombination, challenging established boundaries between sexual and clonal reproduction.

## Results

### Genome assemblies reveal haplotype differentiation and reciprocal translocations

We assembled haplotype-phased chromosome-scale genomes for nine *R. tenuis* accessions (18 haplotypes) sampled across different locations in Brazil (see **Online Methods; Supplementary Table 1 and Supplementary Fig. 1**). Despite its predominantly autogamous mating system, *R. tenuis* exhibits extremely high levels of heterozygosity (>2% in all accessions), exceedingly even that of the closely related outbreeding species *Rhynchospora breviuscula* of approximately 1% (**Supplementary Fig. 2**). Epigenome profiling further revealed a distinctive chromatin organisation of *R. tenuis* holocentromeres. Unlike *Rhynchospora pubera*^17^ and *R. breviuscula*^18^, which feature canonical centromeric domains enriched for CENH3 and CpG DNA methylation, *R. tenuis* holocentromeres display marked CpG hypomethylation and an unexpected enrichment of the active histone mark H3K4me3, coinciding with *Tyba* satellite arrays^19^ (**Extended Data Fig. 1**). These features indicate that *R. tenuis* has evolved an unconventional and atypically euchromatic centromeric environment, potentially linked to its unusual meiotic system and distinctive genome dynamics.

Macrosynteny with closely related species reveals a stepwise fusion trajectory underlying the remarkably reduced chromosome number of *R. tenuis*. Conserved collinear blocks reconstruct an ancestral *Rhynchospora* karyotype of *n* = 5, as seen in *R. breviuscula* and most *Rhynchospora* species with five chromosomes^17,20^. Two successive end-to-end translocations in a common ancestor appear to have produced an intermediate *n* = 3 karyotype by fusing ancestral chromosomes 2 + 5 and 3 + 4, an arrangement retained in the autohexaploid (6*x*) *R. austrobrasiliensis*, the closest known relative of *R. tenuis*, which shares identical fusion breakpoints (**Fig. 1a and Supplementary Fig. 3**). A subsequent fusion involving ancestral chromosome 1 and the 2 + 5 yielded the present *n* = 2 karyotype, conserved across all sequenced *R. tenuis* accessions (**Fig. 1a and Extended Data Fig. 2**).

**Fig. 1.**
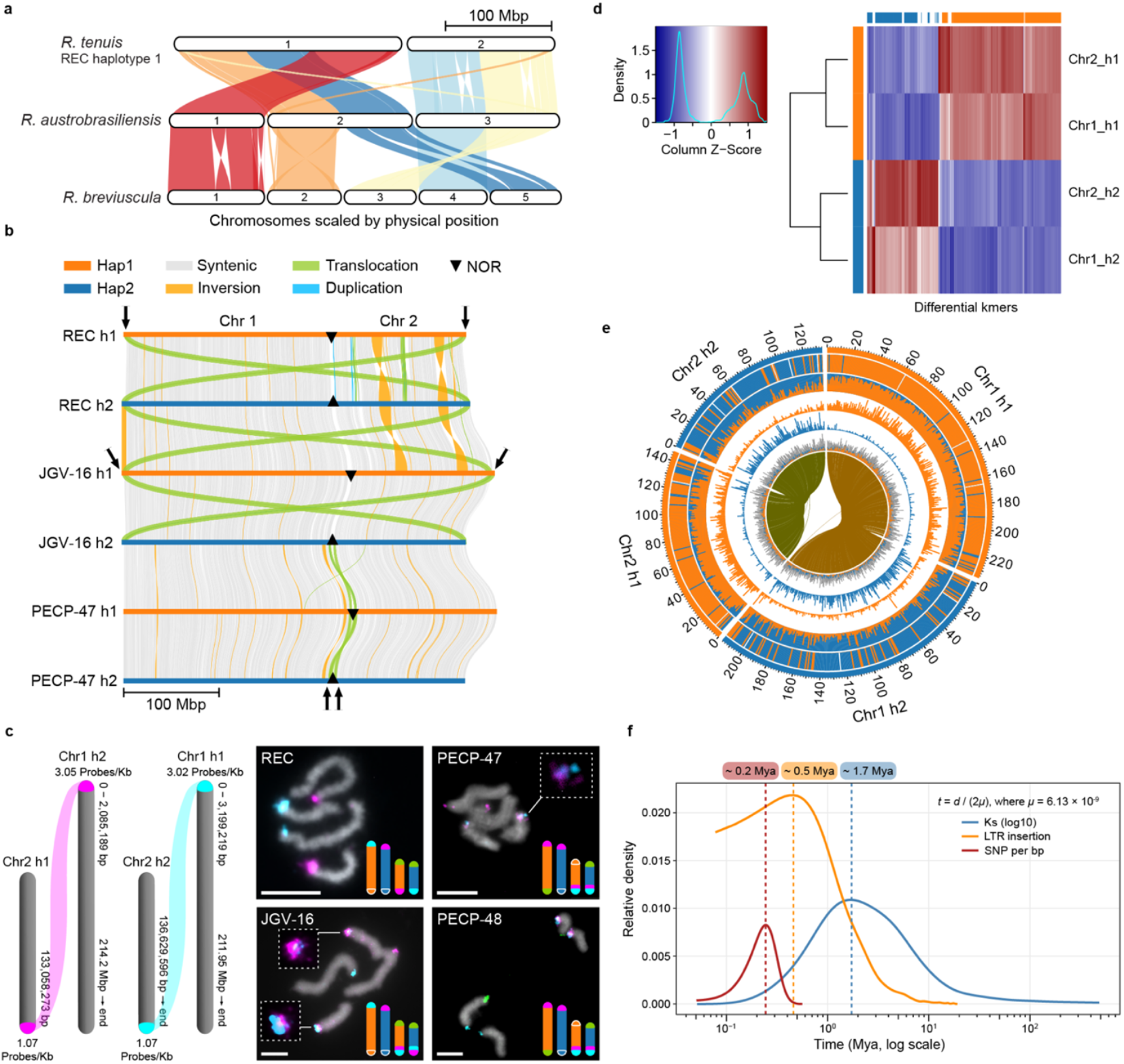
Overview of *R. tenuis* pangenome organisation. (**a**) Macrosynteny across *Rhynchospora* delineates a stepwise reduction in *R. tenuis* chromosome number: conserved syntenic blocks support an inferred ancestral karyotype of *n* = 5, fusion events yielding *n* = 3 in *R. austrobrasiliensis*, and a final fusion producing *n* = 2 in all *R. tenuis* accessions. (**b**) Pangenome analysis also identifies ∼1 Mb terminal reciprocal translocations at one end of each chromosome in *R. tenuis*. (**c**) Representation of the target regions used for haplotype-specific probes design and the final density of each synthesised probe. Oligo-FISH with translocation-specific probe sets (magenta, oligo-probe 1; cyan, oligo-probe 2) validates reciprocal exchanges in the REC and the JGV accessions, but not in PECP-47/PECP-48. 45S rDNA signals (green) in PECP-47/PECP-48 are found on different homologous chromosomes. (**d–e**) *k*-mer-based genome subphasing reports a striking haplotype differentiation found in all *R. tenuis* accessions, which is caused by a differential accumulation of LTR retroelements. (**d**) Unsupervised hierarchical clustering (the horizontal colour bar at the top of the axis indicates to which subgenome the *k*-mer is specific; the vertical colour bar on the left of the axis indicates the subgenome to which the chromosome is assigned). The heatmap indicates the *Z*-scale relative abundance of *k*-mers. The larger the *Z* score is, the greater the relative abundance of a *k*-mer). (**e**) Chromosomal characteristics. From the outer to inner circles (1–9): (1) subgenome assignments based on the *k*-means algorithm; (2) significant enrichment of subgenome-specific *k*-mers − the same colour as the subgenome indicates significant enrichment for those subgenome-specific *k*-mers − white areas are not significantly enriched; (3) normalized proportion (relative) of subgenome-specific *k*-mers; (4–5) count (absolute) of each subgenome-specific *k*-mer set; (6) density of long terminal repeat retrotransposons (LTR-RTs) − if the colour is consistent with the subgenome, it indicates that LTR-RTs are significantly enriched in those subgenome-specific *k*-mers; grey indicates nonspecific LTR-RTs; and (7) homoeologous blocks. All statistics (2–7) are computed in sliding windows of 1 Mb. (**f**) Divergence time estimates between the two non-recombining haplotypes of *R. tenuis* based on the distribution density of pairwise *Ks* (log10) of orthologous coding sequences, haplotype-specific LTRs, and SNP density in the syntenic regions.

In addition to these chromosomal fusions, pangenome-scale synteny uncovered striking chromosome size asymmetry between opposing haplotypes (i.e., h1 and h2), varying from 8 Mbp in the REC (Recife) accession to up to 32 Mbp in JGV (Jaguariaíva) and PECP (Parque Estadual do Cerrado) accessions (**Fig. 1b, Extended Data Fig. 2**). Furthermore, population-specific telomeric reciprocal translocations were detected, ranging from 0.53 to 3 Mbp at one end of each chromosome in *R. tenuis*, with the exchanged segments mapping consistently to different chromosomal ends in different accessions, being classified into three translocation types (**Fig. 1a–b, Extended Data Fig. 2**). To further validate the reciprocal translocations, we designed translocation-specific oligo probes based on the translocation type 1, found in the REC accession (see **Methods**). Fluorescent *in situ* hybridisation (FISH) with translocation-specific oligo- and 45S rDNA (nucleolar organising regions, i.e., NORs) probes validated the reciprocal translocations in all accessions (**Fig. 1c, Extended Data Fig. 3**). The REC accession shows the signals for the translocation oligo-probe 1 (magenta) and oligo-probe 2 (cyan) at the chromosome ends of the non-homologous chromosomes (**Fig. 1b–c, Extended Data Fig. 2 and 3**). JGV accessions showed the same arrangement as REC for chromosome 1, but only a partial translocation was found at the chromosomal ends of chromosome 2 (**Fig. 1b– c, Extended Data Fig. 2 and 3**). As expected from the assemblies, the PECP accessions did not exhibit translocation for that specific chromosome ends. However, reciprocal translocations were found at the other chromosome ends, which in one of the combinations contain the NORs (**Fig. 1b– c, Extended Data Figs. 2 and 3**). No reciprocal translocations were found in the close relative *R. austrobrasiliensis* (2*n* = 6*x* = 18) (**Supplementary Fig. 4**). Our results highlight the role of population-specific structural rearrangements on the unique genome organisation of *R. tenuis*.

Since *R. tenuis* has been reported to be achiasmatic, neither recombination nor segregation of homologous chromosomes is expected, a condition consistent with the ‘Meselson effect’. In such a system, homologous chromosomes accumulate mutations independently, leading to greater divergence between haplotypes than among individuals or populations, effectively generating subgenome-like differentiation in an otherwise diploid genome^11,12^. *K*-mer similarity of all *R. tenuis* haplotypes shows a textbook signature of ‘Meselson effect’, where the same haplotypes from all accessions cluster together while two haplotypes from the same individual do not cluster (**Extended Data Fig. 4**), mimicking an obligately asexual species^12^.

To test whether the *R. tenuis* genome evolves under this regime, we have used subphaser^21^, a *k*-mer-based approach normally used to phase subgenomes in allopolyploid species (polyploid hybrids), on phased haplotypes to estimate divergence between haplotypes versus among-individual divergence. Remarkably, this analysis revealed pronounced subgenome-like differentiation between the two haplotypes across all accessions (**Fig. 1d–e; Extended Data Fig. 5**). This differentiation is mainly explained by haplotype-specific accumulation of TEs, which explains on average 84% of the striking chromosome size asymmetry found between haplotype 1 and 2, while other sequence types, e.g. *Tyba* repeats and genes, did not differ greatly between haplotypes (**Fig. 1d–e; Extended Data Fig. 6**). The extent and distribution of these structural variants, together with population-specific terminal reciprocal translocations, suggest long-term suppression, and effective absence, of meiotic recombination between haplotypes.

Next, we aimed to date the divergence between the two non-recombining haplotypes of *R. tenuis*. Considering that both haplotypes evolve independently without genetic exchange, we estimated their divergence time using synonymous substitution rates (*K*s) calibrated with a fixed molecular clock (6.13e-09 per site per year^22^), and contrasted these values with insertion age estimates of haplotype-specific LTR retrotransposons and SNP-based divergence estimates. The *K*s-based molecular clock suggests that the two haplotypes diverged approximately 1.7 million years ago (Mya; **Fig. 1f**), whereas LTR insertion times indicate a more recent split, between 400,000 and 600,000 years ago (**Fig. 1f**). Divergence times between haplotypes calculated from SNPs divergence were estimated to range from 230,000 - 310,000 years for chromosome 1 and 220,000 - 275,000 years for chromosome 2 (**Fig. 1f**). The higher divergence time inferred from *K*s likely reflects intrinsic differences between coding and non-coding evolutionary rates and the use of a fixed molecular clock, rather than artefactual inflation, whereas the more recent SNP- and LTR-based estimates probably better capture the effective timescale of genome-wide divergence following recombination loss.

### Absence of pairing and synapsis during *R. tenuis* achiasmatic inverted meiosis

Our previous study reported the absence of chiasmata and inverted meiosis in *R. tenuis*^3^. Here, we further tested whether achiasmy and inverted meiosis are consistent features across different populations of *R. tenuis* (REC, PECP and JGV). Indeed, all genotypes analysed always display four univalents at diakinesis and no bivalents (n = 10, n = 37; n = 51), followed by equational division of sister chromatids (n = 37, n = 28; n = 12), consistent with the concept of inverted meiosis; (**Fig 2a–d; Supplementary Fig. 5**). To deepen our understanding, we further tracked the localization of translocation-specific oligo- and 45S rDNA probes on meiotic chromosomes. During early meiotic prophase, translocation-specific oligo-probes and telomeric signals are never associated indicating absence of pairing and synapsis, while rDNA regions are associated at the nucleolus (n = 55; **Fig. 2a–b; Supplementary Fig. 6a**), followed by diakinesis with four univalents and no bivalents, indicating asynapsis and achiasmy (**Supplementary Fig. 6b**).

**Fig. 2.**
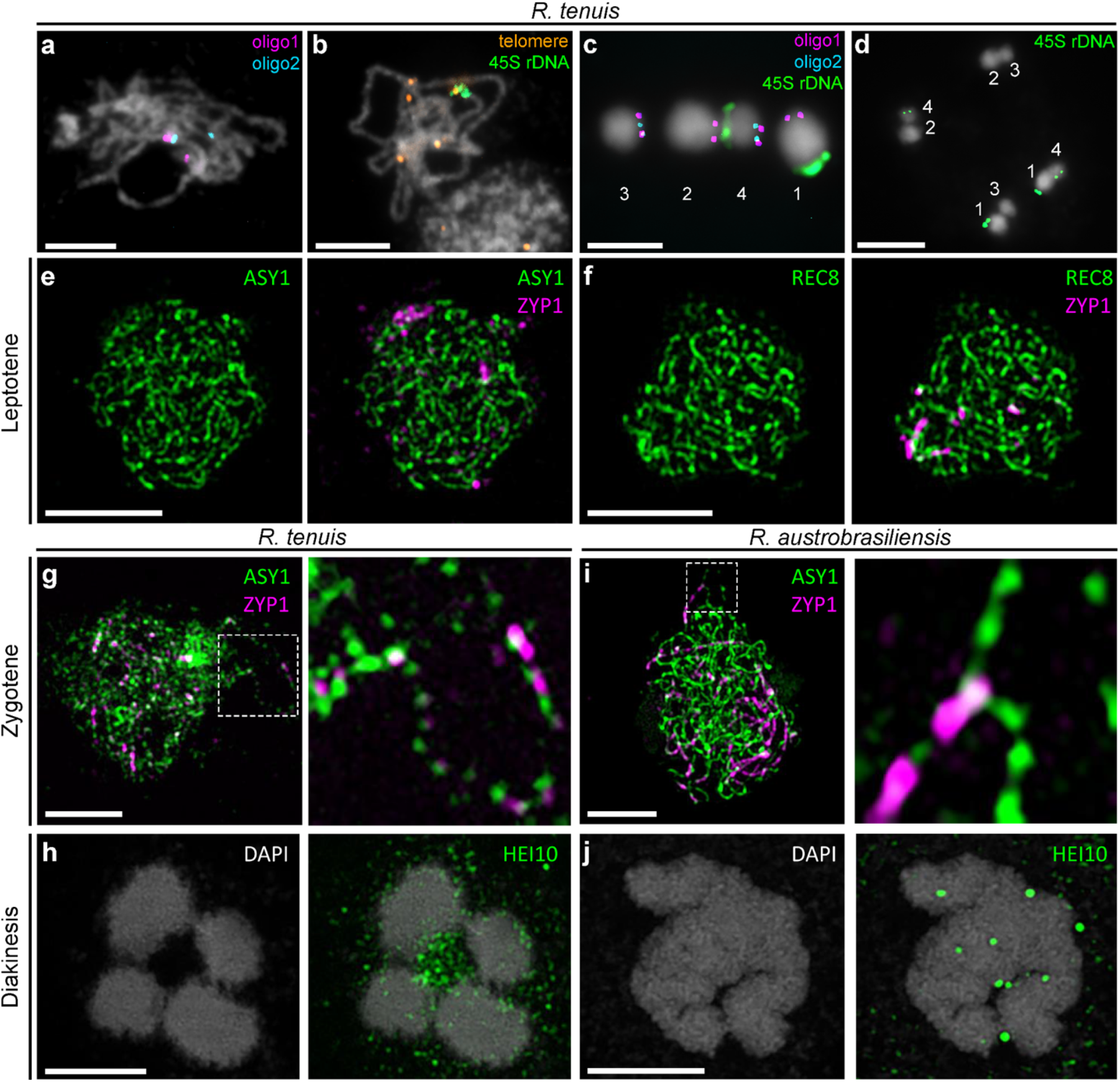
FISH and Immunostaining confirm the failure of pairing, synapsis and crossover formation in *R. tenuis* (REC). FISH using (**a**) translocation-specific probe sets and (**b**) 45S rDNA and telomeric probes revealed neither pairing of translocation regions (**a**) nor bouquet formation (**b**) during early meiosis. FISH with translocation-specific probes (**c**) and 45S rDNA (**c–d**) in metaphase I (**c**) and telophase II (**d**). 1 = Chr1_h1, 2 = Chr1_h2, 3 = Chr2_h1 and 4 = Chr2_h2. Immunostaining with (**e**) ASY1 and (**f**) REC8 together with ZYP1 signal in Leptotene. Immunostaining with (**g; i**) ASY1 and ZYP1 depicts abnormal zygotene in *R. tenuis* (**g**) contrasted to zygotene in *R. austrobrasiliensis*, which shows normal axis and synapsis (**i**). (**h; j**) HEI10 immunostaining shows the lack of foci at diakinesis in *R. tenuis* (**h**), but the presence of foci in *R. austrobrasiliensis* (**j**). Scale bars correspond to 5 μm.

At metaphase I, univalents aligned perpendicular to the equatorial plate and sister chromatids exhibited bi-orientation, leading to segregation of sisters at anaphase I (equational division) and formation of two diploid products at telophase I/interkinesis (**Fig. 2c; Supplementary Fig. 6c**). During meiosis II, the previously separated sisters realigned as homologous non-sisters at metaphase II and segregated at anaphase II, yielding four haploid products (**Fig. 2d; Supplementary Fig. 6d**). Across all populations, the consistent absence of chiasmata/bivalents and the inversion of the canonical meiotic order (equational division in meiosis I and reductional division in meiosis II) demonstrate that achiasmy and inverted meiosis are conserved and consistent features of *R. tenuis* rather than accession-specific anomalies.

We extended our analysis of meiotic progression in *R. tenuis* by immunostaining male meiocytes with antibodies against the axis components ASY1^23^ and REC8^24^, the synaptonemal complex (SC) transverse filament ZYP1^25^, and the CO marker HEI10^26^. Early prophase I appeared conserved: ASY1 (n = 41) and REC8 (n = 29) localised as linear signals along chromosomal axes in all cells analysed (**Fig. 2e–f and Extended Data Fig. 7a–b**). By contrast, ZYP1 failed to assemble as a continuous SC; instead, it formed a few persistent short elongated structures and polycomplexes while ASY1 was unloaded from the axes (n = 10; **Fig 2g and Extended Data Fig. 7c**). HEI10 was occasionally detected on ZYP1 polycomplexes but did not form discrete foci along chromosome axes (n = 12; **Extended Data Fig. 7d**). At diakinesis, meiocytes invariably displayed four univalents, and no HEI10 foci indicative of COs were observed (n = 21; **Fig 2h**). These features, fragmented SC signal, and absence of CO markers, culminated in univalents at metaphase I (**Fig. 2c and h**), consistent with achiasmy.

As a contrast, the closest relative, *R. austrobrasiliensis*^27^, exhibited canonical chiasmatic meiosis. ASY1 localized to the axes and was progressively unloaded during synapsis (n = 12), the SC was assembled as continuous ZYP1 signals (n = 12; **Fig 2i and Extended Data Fig. 8a**), and HEI10 initially appeared as closely spaced foci on ZYP1-marked regions, later re-localising as a few intense foci on bivalents, co-localizing with MLH1 (n = 59) and confirming normal CO maturation (**Fig. 2j and Extended Data Fig. 8b–d**). To explore a genetic basis for recombination failure in *R. tenuis*, we surveyed meiotic gene content and transcription. Most core meiotic genes and their corresponding transcripts are present (**Supplementary Table 2**). However, we detected haplotype-specific copy-number variation at several meiotic loci, including additional copies for *ASY1* and *MSH5* specific to achiasmatic *R. tenuis* (**Supplementary Table 2**). *SHOC1*, responsible for the maturation of mid-to-late recombination intermediates in plants and mammals^28,29^, showed some ambiguities. Two independent RNA-seq datasets did not contain traces of *SHOC1* expression. However, the gene displays a complete and apparently functional sequence, and PCR amplification of the functional domain from tissue-specific cDNA was successfully achieved. However, we cannot rule out that expression levels or post-transcriptional modifications might affect the activity of the SHOC1 protein.

### Genetic COs are absent in both male and female meiosis

To test genetically whether COs are absent in *R. tenuis*, we applied our single-gamete sequencing pipeline^18,20^ to pollen nuclei across five geographically distinct accessions, leveraging chromosome-scale, haplotype-phased assemblies as references. Using high-throughput single-nucleus profiling (snRNA- and snATAC-seq), we obtained 10,997 pollen grains in total that yielded high-confidence, genome-wide genotypes (**Fig. 3a, Supplementary Table 3**). Haplotype blocks spanned entire chromosomes with no confident COs detected in any gamete, demonstrating that male achiasmy is pervasive across accessions. This means that, if crossovers occur at all, they must be extremely rare, consistent with a complete suppression of recombination during male meiosis. Because our single-pollen sequencing interrogates only male meiosis, we assessed female recombination genetically by sequencing the offspring of controlled self-crossing heterozygous mother plants. In the highly heterozygous genome of *R. tenuis*, female COs would be visible as haplotype switch points in F1 individuals. Whole-genome sequencing of 49 and 62 F1 individuals coming from the controlled self-crossed seeds from heterozygous mothers of REC and PECP accessions, respectively, revealed no recombination breakpoints on either chromosome in any offspring (**Fig. 3b**). Remarkably, all F1 individuals carried the same two intact parental haplotypes as the mother, with no detectable gene conversion, demonstrating that female meiosis, like male meiosis, is achiasmatic. Furthermore, these results also indicate that all offspring show clonal-like formation as they mirror the maternal genotype.

**Fig. 3.**
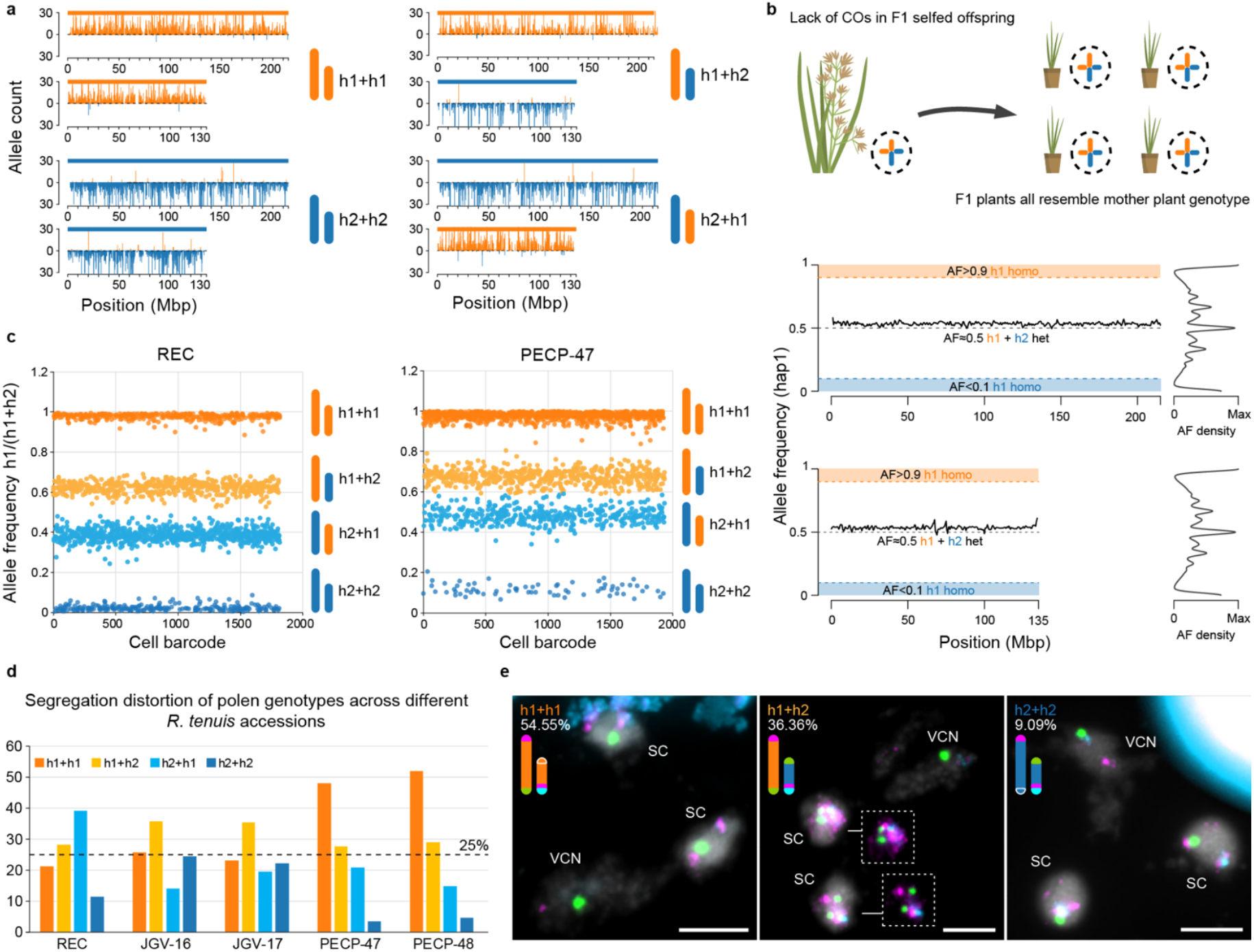
Single-gamete sequencing reveals complete suppression of crossovers and male meiotic drive under asymmetric microsporogenesis in *R. tenuis*. (**a**) Four examples of pollen genotype combinations obtained from single-nucleus sequencing of pollen nuclei show chromosome-scale haplotype blocks with no recombinants. Failing to detect a single CO after analysing 10,997 pollen nuclei from five different accessions. (**b**) F1 offspring of controlled selfing-crossed mother plants show consistent chromosome-wide 1:1 allele ratios, indicating the absence of COs and mirroring of the mother genotype, indicating clonal-like seed formation. (**c**) Allele frequency, as defined by the ratio of h1 markers divided by the total markers (h1+h2) identified in each single pollen nucleus sequenced, reveals four main groups of pollen genotypes in *R. tenuis*. (**d**) Genotype class frequencies highlight segregation distortion among sequenced pollen genotypes across accessions, with the strongest bias in PECP-47 and PECP-48 favouring the larger, TE-rich haplotype 1. (**e**) Representative haplotype-specific barcoding across chromosomes illustrates intact, nonrecombinant transmission in individual mature pollen grains. In PECP-47/PECP-48, reciprocal translocations of the NOR eliminate NOR-null gametes post-meiotically, further skewing haplotype transmission. SC = sperm cell, VCN = vegetative cell nucleus. Scale bar = 5 µm. Oligo-probe 1 = magenta, oligo-probe 2 = cyan, 45S rDNA probe = green.

We next examined transmission patterns among *R. tenuis* viable pollen grains, which are lower than 75% across all accessions analysed (**Supplementary Fig. 7**). In *R. tenuis*, as a conserved feature of the *Rhynchospora* genus, male microsporogenesis is asymmetric: only one of the four meiotic products matures into a functional microspore (pollen), while the remaining products degenerate^30–33^ (**Extended Data Fig. 9a**). Such asymmetry provides a cellular context for male meiotic drive, which is not expected under symmetric microsporogenesis. Consistent with this, allelic counts at haplotype-informative markers revealed strong, accession-specific segregation distortion among viable pollen (**Fig. 3c**). Under inverted meiosis, sister chromatids segregate at meiosis I, yielding diploid secondary spermatocytes; if meiosis II followed Mendelian segregation and all four products matured equivalently, the four genotype classes defined by the two chromosomes should be equally frequent. Instead, in 10,997 pollen nuclei with complete genotype calls across both chromosomes, we observed significant over-representation of specific chromosome combinations.

Across the full dataset, distortion was particularly substantial in two closely related accessions (PECP-47, PECP-48), where the larger, TE-rich haplotype 1 was preferentially transmitted in mature pollen (>50%) over other combinations (**Fig. 3d**). Remarkably, pollen cells with both chromosomes of haplotype 2 were found at very low frequency (4%). Given that haplotype 1 exceeds haplotype 2 by ∼32 Mb primarily due to LTR retrotransposon accumulation, these results point to a male meiotic drive system in which TE-associated features may directly or indirectly skew chromosomal inheritance during, or immediately following, meiosis II. Together with the complete absence of COs, these findings indicate that, in the asymmetric microsporogenesis of *R. tenuis*, transmission bias operates in the context of recombination loss to favour particular haplotype combinations, with chromosome size being the strongest factor.

A structural basis for this distortion emerges from the position of the nucleolar organiser region (NOR, 45S rDNA). In REC and JGV, the NOR is not interchanged between non-homologous chromosomes, whereas in PECP-47 and PECP-48, the NOR-bearing segments are reciprocally translocated between Chr1 and Chr2. Consequently, one genotype class, Chr1_h2 + Chr2_h1, lacks any NOR, predicting compromised ribosome biogenesis and gamete inviability. Intriguingly, single-nucleus genotypes retrieved pollen grains in PECP-47/PECP-48 showing the following frequencies: Chr1_h1 + Chr2_h1 (50%, bearing one NOR), Chr1_h1 + Chr2_h2 (28%, bearing two NORs), Chr1_h2 + Chr2_h1 (18%, NOR-null), and Chr1_h2 + Chr2_h2 (4%, bearing one NOR). However, FISH on PECP-48 mature pollen grains (n = 33), using a combination of oligo-probes and 45S rDNA, recovered similar abundances as detected by our single-pollen sequencing, but only on genotype combinations that retain at least one NOR. However, we never found the NOR-null class (Chr1_h2 + Chr2_h1) genotype combination in mature pollen grains and in sperm cells from germinated pollen tubes (**Fig. 3d–e and Extended Data Fig. 9b**). Remarkably, we found in 90.5% (n = 21) of the germinated pollen tubes only one single NOR signal in both tube nucleus and sperm cells, which is consistent with chromosomes from the same haplotype being inherited together (**Extended Data Fig. 9b**). Further investigation in earlier pollen stages consistently revealed segregation distortion of haploid products, with the NOR-null class genotype being found only at the degenerative cells (non-selected products) during pollen development (**Extended Data Fig. 9c–d**), indicating post-meiotic selection of haploid products during early pollen development. Together with a consistent bias favouring the larger, TE-rich haplotype 1 in PECP-47/PECP-48, these results reveal asymmetric male meiotic drive in *R. tenuis*, in which structural features, including NOR placement and TE-associated genome expansion, skew chromosomal transmission in the complete absence of recombination.

### Post-meiotic selection yields clonal-like offspring despite sexual reproduction

To exclude that the obtained seeds from our F1 offspring are derived from apomixis, a form of asexual seed formation in which embryos develop without fertilisation or meiosis^34^, we performed crossing experiments to confirm that seed formation requires fertilisation. Remarkably, viable seeds were only obtained when both parents carried the same reciprocal translocation type (e.g., PECP-47 × PECP-48), whereas all other combinations failed to set seed (**Fig. 4a**). Sequencing of offspring from these crosses confirmed that all plants were heterozygous, consistent with fertilisation but indicating that heterozygosity is obligatory (**Fig. 4b; Supplementary Dataset 1**).

To further examine female gametophyte and early seed development, we performed histological analyses of early and mature flowers. Female gametophytes showed normal development, with well-differentiated embryo sacs and initiation of seed development consistent with fertilisation (**Fig. 4c**). At early stages of megagametogenesis, multiple embryo sacs were frequently observed out of which only one continues to mature in 98% of the cases (n = 59, **Supplementary Fig. 8**). Pollen tube growth towards the ovule was consistently observed (**Fig. 4d and Extended Data Fig. 10a–b; Supplementary Fig. 9 and 10**). Although healthy embryos were consistently observed developing from the micropyle, indicating sexual origin (**Fig. 4e–f**), a high proportion of ovules seem to fail to develop into seeds (**Fig. 4h and Extended Data Fig. 10c–f**). Microscopic inspection of these undeveloped ovules revealed intact, likely unfertilised gametophytes, suggesting failure of fertilisation, potentially due to male gametophytic defects, whereas others likely collapsed shortly after fertilisation (**Fig. 4h and Extended Data Fig. 10d–f**). Furthermore, a high frequency (87%) of aborted seeds was observed (n = 167, 145 aborted: 22 fertile for PECP-48; **Extended Data Fig. 10d–f**), indicating strong post-meiotic and post-fertilisation selection.

**Fig. 4.**
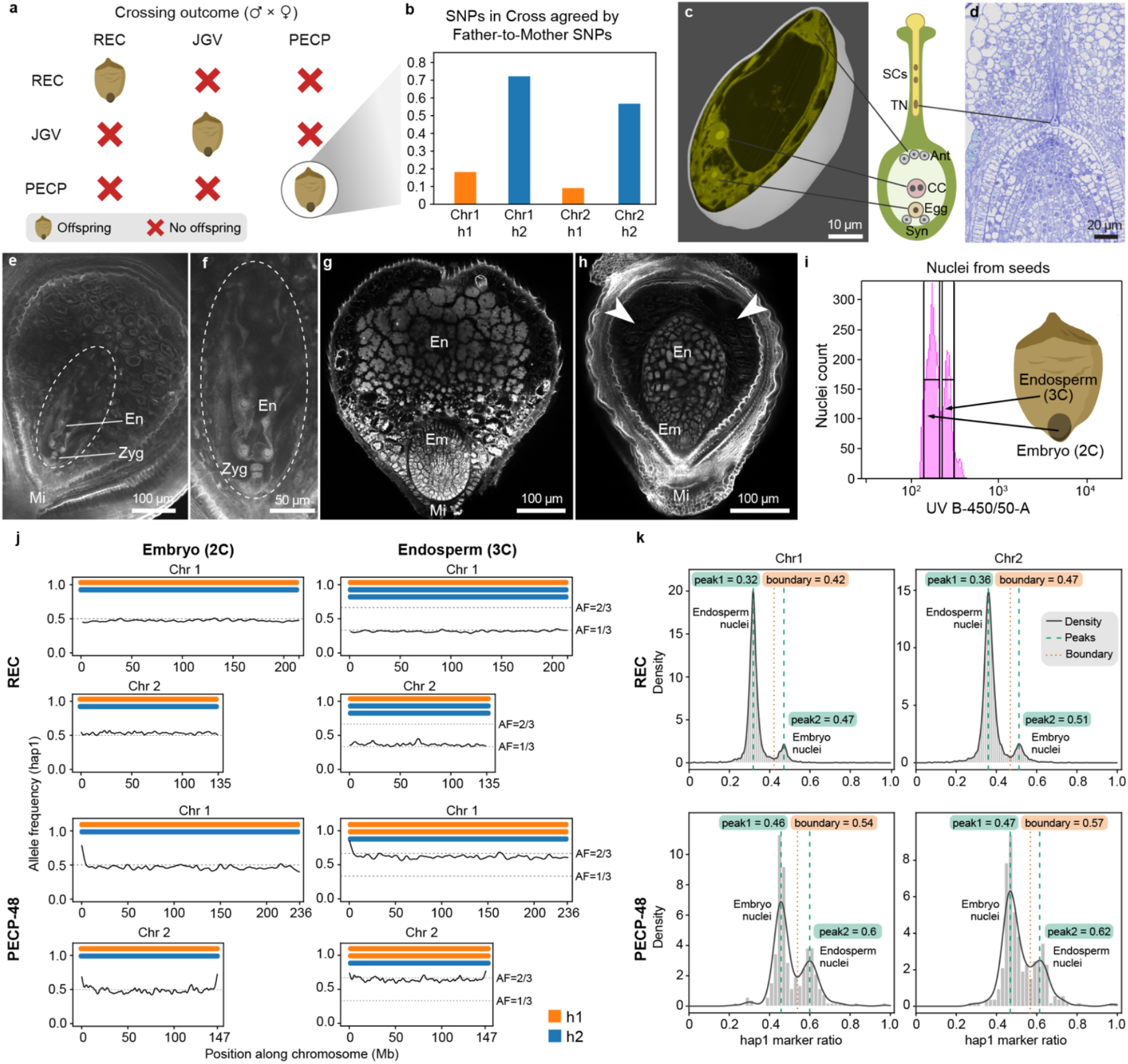
Female achiasmy and post-meiotic selection produce clonal-like progeny despite sexual reproduction. (**a**) Crossing experiments between accessions with different reciprocal translocation genotypes show seed formation only when both parents share the same translocation type (example: PECP-48 × PECP-47). All successful crosses yield heterozygous offspring. (**b**) In the cross PECP-48 × PECP-47, alignment of progeny reads to the maternal genome (PECP-48) shows paternal-specific alleles across haplotype 2, indicating biparental fertilisation with haplotype 1 maternally inherited and haplotype 2 paternally inherited. (**c**) 3D reconstruction of serial histological sections reveals canonical female gametophyte development with a normal embryo sac containing egg cell (Egg), synergids (Syn), central cell (CC) with fused polar nuclei, and antipodals (Ant). (**d**) Pollen tube growth towards the ovule. (**e**) Very young stage after fertilisation showing the Zygote (Zyg) and coenocytic endosperm (En). (**f**) Enlargement of selected region shown in **e**. (**g**) Fully mature seed showing an embryo in a late stage of development surrounded by the endosperm. Embryos were consistently observed developing from the micropyle (Mi) consistent with sexual origin. (**h**) Late aborted seed showing underdeveloped and arrested embryo and endosperm. Arrowheads point to collapsed area. (**i**) Flow cytometry profiles of seed nuclei reveal diploid embryos and triploid endosperms, consistent with canonical double fertilisation and the formation of seeds through sexual reproduction. (**j**) Single-nucleus sequencing of embryo and endosperm nuclei demonstrates biparental contributions: embryos are uniformly heterozygous, while endosperms exhibit the expected 2:1 maternal:paternal dosage. In REC and PECP-48, maternal transmission is biased toward the larger, TE-rich haplotype (haplotype 2 in REC, haplotype 1 in PECP-48). (**k**) Allelic ratio distributions confirm consistent dosage bias and post-meiotic selection that together restore heterozygosity and produce clonal-like offspring despite sexual reproduction.

Flow cytometry of full mature seeds confirmed diploid embryos and triploid endosperms (**Fig. 4i**), consistent with meiosis followed by double fertilisation. Single-nucleus ATAC-sequencing of flow-sorted embryo and endosperm nuclei verified biparental contributions: embryos were uniformly heterozygous, while endosperms showed the expected 2:1 maternal:paternal dosage (**Fig. 4j**), enabling direct inference of female gamete haplotype composition. Strikingly, in both REC and PECP-48 accessions, a single maternal haplotype was always transmitted (haplotype 2 in REC and haplotype 1 in PECP-48; **Fig. 4k**). In both cases, the driving haplotype corresponded to the physically larger, TE-enriched chromosomes.

Importantly, none of the imaged ovules or seeds showed evidence of apomictic development. In all cases, embryos originated exclusively from the micropyle, never from sporophytic tissues, and only a single embryo was observed per seed. Flow cytometry of mature seeds confirmed the expected 2:3 embryo:endosperm ratio, consistent with double fertilisation. These results exclude diplospory, apospory, and adventitious embryony as mechanisms of seed formation, supporting sexual reproduction as the sole reproductive mode in *R. tenuis*.

Our findings indicate that seed production in *R. tenuis* is strictly sexual, but its success depends on translocation compatibility and the maintenance of heterozygosity, which appears to be essential for embryo viability. Together with the single-gamete sequencing data, these findings establish obligate, genome-wide achiasmy in both sexes. The apparent paradox of clonal-like offspring arising through sexual reproduction can be explained by extreme segregation distortion, reinforced by post-meiotic selection, which ensures that only heterozygous zygotes with specific haplotype combinations survive. As a result, viable progeny genetically mirrors the maternal genotype while retaining biparental genomic contributions (**Fig. 5**).

**Fig. 5.**
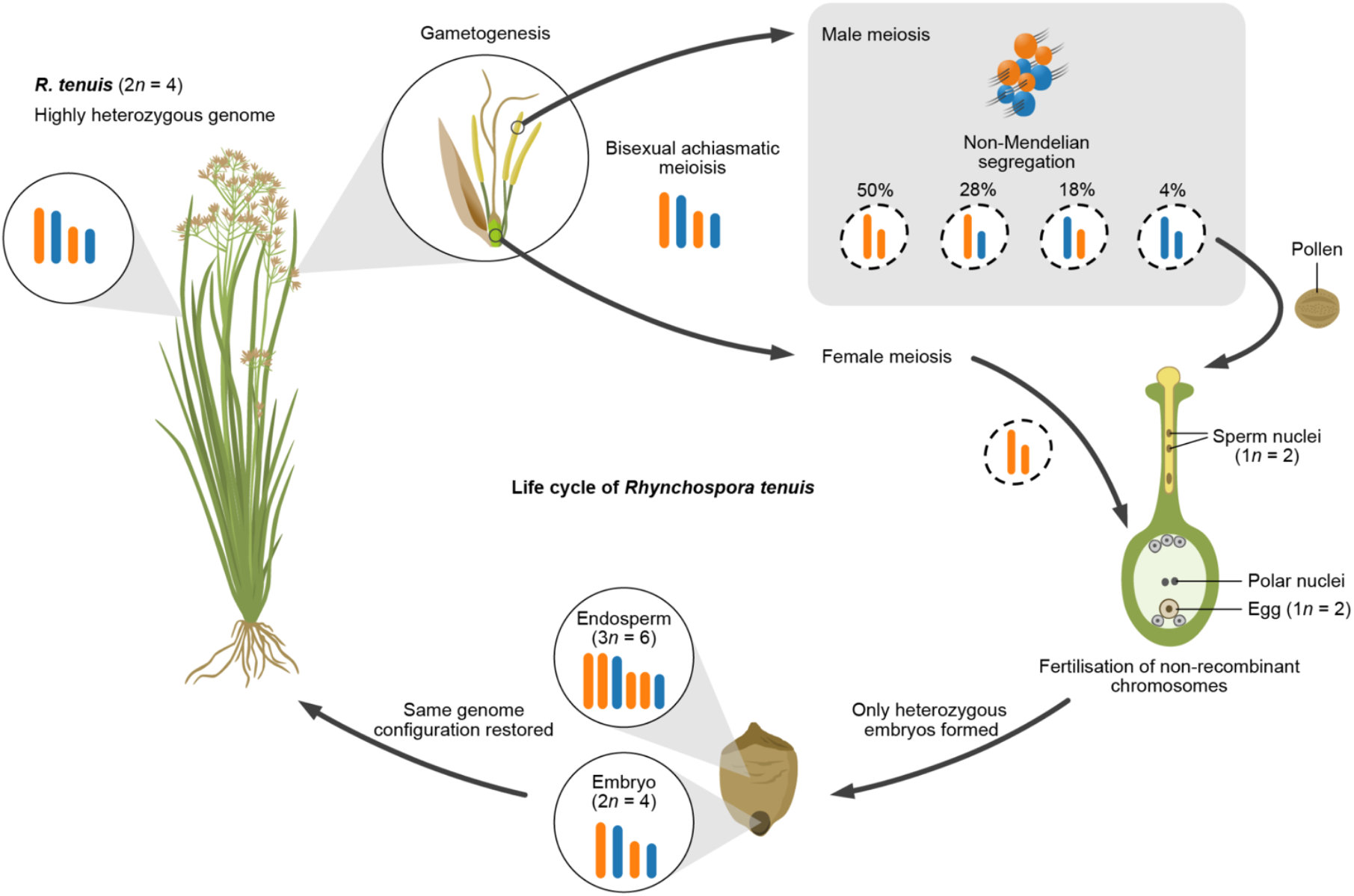
Life cycle of *Rhynchospora tenuis* (i.e. PECP-48 genotype) under achiasmatic meiosis. Schematic representation of the *R. tenuis* life cycle (2*n* = 4). Both male and female meiosis occur without crossovers (achiasmatic inverted meiosis). Male gametogenesis is asymmetric, yielding a single functional sperm nucleus and non-Mendelian segregation. Fertilisation restores the diploid state with intact haplotypes, but only heterozygous seeds develop into viable embryos, ensuring faithful restoration of the highly heterozygous parental genome each generation.

## Discussion

Our study identifies a reproductive system without precedent: a sexually reproducing plant that has completely lost meiotic recombination while maintaining accurate chromosome transmission and normal fertility. In *R. tenuis*, we demonstrate obligate achiasmy in both sexes, characterised by the absence of crossovers and gene conversion, despite the normal initiation of meiotic processes. Thousands of sequenced gametes and multiple progenies reveal intact parental haplotypes and offspring genetically identical to their heterozygous mothers. This clonal-like inheritance arises through extreme segregation distortion, haplotype-biased transmission, and strong post-meiotic selection that restricts viable zygotes to specific heterozygous combinations.

These findings challenge the view that meiotic recombination is essential for proper chromosome segregation and for the generation of genetic diversity^35,36^. Although achiasmy occurs in limited contexts, such as *Drosophila* males^7^, *Bombyx mori* females^10^, or non-recombining sex chromosomes^2^, *R. tenuis* appears to represent the first case of complete, genome-wide achiasmy in both sexes of an otherwise sexually reproducing multicellular eukaryote. A partial parallel exists in the budding yeast *Saccharomycodes ludwigii*, which undergoes meiosis with extreme suppression of CO while retaining SPO11-dependent double-strand break formation and high heterozygosity through intratetrad mating^37^. However, unlike *R. tenuis*, this system operates in a unicellular fungus with automixis and lacks the developmental, cytological, and selective constraints imposed by multicellular reproduction, gametophyte competition, and post-zygotic viability. Moreover, the unusual karyotype of *R. tenuis* provides a permissive framework, as holocentric chromosomes stabilise segregation in the presence of rearrangements^17^, inverted meiosis reduces reliance on chiasmata^3^, and an extremely reduced chromosome number minimises the risk of missegregation. Together, these features create a cellular context in which recombination can be lost without catastrophic consequences.

The system both parallels and diverges from other non-Mendelian inheritance mechanisms. In *Ooceraea biroi* ants and *Mesorhabditis belari* nematodes, heterozygosity is retained through co-segregation of recombinant chromatids even when recombination is active^38,39^. A related plant analogue occurs in *Oenothera*, where balanced lethal and extensive reciprocal translocations enforce permanent translocation heterozygosity and restrict recombination to subterminal regions, thereby maintaining heterozygous haplotypes across generations^40–43^. By contrast, *R. tenuis* achieves the same functional outcome through a combination of bisexual achiasmy and gametic selection.

At first sight, clonal-like inheritance and recombination loss resemble the Meselson effect, which predicts that haplotypes in asexual lineages diverge through independent mutation accumulation^11,12^. Indeed, *R. tenuis* exhibits functional asexuality, and haplotypes are structurally divergent.

However, unlike the parthenogenetic booklouse *Liposcelis bostrychophila*, where genomic data support Meselson-driven haplotype divergence^44^, *R. tenuis* avoids this outcome. Meiosis and fertilisation are retained, homozygotes are eliminated, and meiotic drive biases transmission toward TE-rich haplotypes. These features limit independent haplotype evolution and reduce the risk of long-term genomic degeneration.

Recent work in the planarian *Schmidtea mediterranea* reveals that stepwise structural rearrangements can progressively suppress recombination before the evolution of fissiparous asexuality^45^. Both species show extensive structural variation that remodels the recombination landscape and produces divergent haplotypes without immediate genomic collapse. However, the evolutionary trajectories differ. In planarians, recombination suppression is followed by the loss of sex and reliance on somatic selection to mitigate mutation load. In *R. tenuis*, recombination disappears mechanistically through obligate achiasmy, but meiosis and fertilisation persist. Clonal-like inheritance resulting from meiotic drive and post-meiotic selection rather than a shift to asexual reproduction. This highlights how similar ingredients, including recombination suppression and transmission bias, can generate different resolutions of the tension between sex and clonality.

The basis of recombination failure in *R. tenuis* remains unclear. Although *ASY1* and *MSH5* are duplicated, early meiotic processes appear to initiate normally before pairing and synapsis fail. Structural divergence between haplotypes, including reciprocal telomeric translocations and asymmetric TE accumulation, may disrupt homologous recognition. We observe consistent female drive of a single haplotype in independent accessions, in each case favouring the TE-rich, larger chromosomes. Because kinetochore activity extends along the entire chromosome in holocentric chromosomes during *Rhynchospora* meiosis^3,18,33^, TE accumulation could enlarge the functional kinetochore and bias segregation in line with centromere and holokinetic drive models^46–48^.

TE expansion on a single haplotype may reflect multiple processes. Germline-specific TE mobilisation during gametophyte development^49,50^, could gradually enrich the haplotype that is preferentially transmitted through the ovule. Alternatively, haplotype asymmetry may derive from ancient hybridisation followed by recombination suppression. The opposite direction of female drive in different accessions (haplotype 1 in PECP and haplotype 2 in REC) argues that haplotype bias is dynamic rather than fixed, possibly reflecting local selection or stochastic changes in drive polarity.

Obligate bisexual achiasmy in *R. tenuis* expands the known limits of meiotic flexibility and blurs the distinction between sexual and clonal reproduction. Although recombination is widely regarded as essential for avoiding mutational meltdown and promoting adaptive potential^14,51–53^, our results show that clonal-like inheritance can arise through modifications to meiotic mechanics, genome architecture, and post-meiotic selection without apomixis. Whether similar systems exist in other holocentric lineages with very low chromosome numbers remains unknown.

In conclusion, *R. tenuis* demonstrates that meiosis and fertilisation can persist in the complete absence of COs, creating a reproductive system that bridges sex and clonality. Holocentricity, inverted meiosis, and meiotic drive combine to stabilise obligate achiasmy. Our findings establish *R tenuis* as a powerful model for studying how recombination can be lost, replaced, or reconfigured, providing a new framework for investigating the evolutionary consequences of extreme meiotic adaptations.

## Methods

### Plant material

Plants of *R. tenuis* and *R. austrobrasiliensis* were collected from natural populations in Brazil under appropriate permits and cultivated under controlled greenhouse conditions. Nine geographically distinct accessions of *R. tenuis* were used for genome sequencing and cytological analyses. Individual flowers and pollen were staged for meiotic analyses. All experimental material was propagated clonally from field-collected individuals to ensure genetic identity across assays.

### Sequencing

High-molecular-weight DNA was extracted from young leaf tissue using a modified CTAB protocol. PacBio HiFi libraries were prepared and sequenced on the Sequel IIe platform. Illumina short-read sequencing was used for genome polishing and transcriptome profiling. Hi-C libraries were constructed using Arima Hi-C kits and sequenced on an Illumina NovaSeq 6000 platform. RNA was extracted from young inflorescences for expression analyses.

### DNA isolation

High-molecular-weight DNA from *R. tenuis* and *R. austrobrasiliensis* was isolated from 1.5 g of material with a NucleoBond HMW DNA kit (Macherey Nagel). Quality was assessed with a FEMTO-pulse device (Agilent), and quantity was measured with a Quantus fluorometer (Promega).

### PacBio

HiFi libraries were prepared according to the “*Preparing whole genome and metagenome libraries using SMRTbell® prep kit 3*.*0*” manual, with an initial DNA fragmentation by Megaruptor-3 (Diagenode) and final size-selection by BluePippin (Sage Science). Size distribution was again controlled by FEMTO-pulse (Agilent). Size-selected libraries were then sequenced on a Revio device with Revio polymerase kit and Revio chemistry for 30 h (*Pacific Biosciences*).

### Arima Hi-C

Plant tissues were cross-linked with 1% formaldehyde for 30 minutes at room temperature, and the reaction was quenched with 125 mM glycine for 10 minutes. Subsequently, the tissues were ground using a TissueLyser at a frequency of 30 Hz for 3 minutes. Nuclei extraction was performed using the CelLytic PN Plant Nuclei Isolation/Extraction Kit (Sigma-Aldrich, Burlington, MA, USA) according to the manufacturer’s protocol. Hi-C libraries were prepared using the Arima High Coverage Hi-C Kit (Arima Genomics, A410110, Carlsbad, CA, USA) following the manufacturer’s instructions, and were then sequenced paired-end (2 × 150 bp) on a NextSeq 2000 instrument (Illumina, San Diego, CA, USA).

### Methyl-seq

To investigate the methylome space in *R. tenuis* (REC accession), the relatively non-destructive NEBNext Enzymatic Methyl-seq Kit was employed to prepare an Illumina-compatible library, followed by paired-end sequencing (2 × 150 bp) on a NextSeq2000 (Illumina) instrument. For each library, 10 Gb of reads was generated.

### RNAseq

Total RNA was isolated from root, leaves, and flower buds (REC accession). Poly-A RNA was enriched from 1 μg total RNA using the NEBNext® Poly(A) mRNA Magnetic Isolation Module. RNAseq libraries were prepared as described in the NEBNext Ultra™ II Directional RNA Library Prep Kit for Illumina (New England Biolabs). A total of 11 cycles were applied to enrich library concentration. Sequencing was performed at BGI Genomics (Hong Kong) using a BGISEQ-500 system on the DNBseq platform in paired-end mode with a read length of 2 x 150 bp.

### Illumina libraries (TPase and DNA FS)

Genomic DNA from 49 and 59 controlled self-crossed *R. tenuis* REC and PECP35-7 accessions, respectively, were deep-sequenced with an Illumina HiSeq 3000 or, alternatively, with DNBseq short read sequencing (BGI Genomics, Hong Kong) in 150-bp paired-end mode.

### Heterozygosity estimation

Heterozygosity levels of *R. tenuis, R. breviuscula*, and all *R. tenuis* accessions were estimated by GenomeScope2.0^54^. K-mer counting was performed using FASTK (v1.1) (https://github.com/thegenemyers/FASTK) with PacBio HiFi reads, which were used for genome assembly, as the input. Kmer size is 31 (Fastk -k31). The resulting binary output (.hist) from FASTK was then converted to a text histogram file by Histex (part of FASTK): Histex -G hifi_31mer.hist > hifi_31mer.histo. The histogram was then analysed with GenomeScope2 using k=31 and ploidy=2, except for *R. austrobrasiliensis*, where ploidy=6.

### Genome assemblies

HiFi reads were assembled using Hifiasm v0.16.1 with default settings that combine both PAcBio HiFi reads and Hi-C reads to generate haplotype-resolved contigs. Assembly quality was assessed using BUSCO v5.2.2^55^ and QUAST^56^. Duplicated haplotypes were retained for downstream analysis.

### Scaffolding

Chromosome-scale scaffolding was performed using the Juicer and 3D-DNA pipelines^57,58^. Contigs from each haplotype generated from HiFiasm were taken as Hi-C alignment targets. Hi-C contact maps were visualised using Juicebox Assembly Tools to manually curate misjoins and validate the final pseudomolecules. Haplotypes were scaffolded independently and phased using *k*-mer-based alignment strategies.

### Gene-based synteny analysis

Genes used for synteny analysis were annotated by a deep-learning-based tool Helixer (v0.3.4)^59,60^ with the land plant mode. Gene synteny was analysed for *R. breviuscula* haplotype 1, *R. austrobrasiliensis* haplotype 1, and all haplotypes of nine *R. tenuis* accessions using GENESPACE (v1.3.1)^61^ running in R (v4.2.0) along with the dependent tools OrthorFinder (v2.5.5)^62^ and MCScanX (v1.0.0)^63^. The R script for running GENESPACE (run_genespace.R) can be found in our project GitHub page: https://github.com/Raina-M/Rhynchospora_tenuis_project.

### DNA sequence-based synteny analysis

Collinearity of all *R. tenuis* haplotypes was analysed by SyRI (v1.5.3)^64^. The alignment between haplotypes was conducted using minimap2 (v2.28)^65,66^ with the following parameter settings: -ax asm5 --eqx. The plot was done by plotsr (v0.5.3) with a minimal 50kbp syntenic block size. NOR regions labelled on the haplotypes were searched by BLAST (v2.12.0) with the *Arabidopsis* rDNA sequences as the template.

### Pangenome analysis

Pangenomes were generated using minigraph v0.17 with phased chromosome-scale assemblies from nine accessions. Variants were extracted using vg toolkit v1.40.0. Structural variants and sequence divergence were quantified across homologous haplotypes. Repeat annotation was performed with DANTE^67^ and EDTA v2.0^68^ using the plant TE library. Pairwise divergence and TE composition were visualised using custom R scripts.

### ChIP-seq

ChIP was performed as described by Reimer and Turck^69^ with minor modifications. Young leaves and flower buds were collected from greenhouse-grown plants and immediately frozen in liquid nitrogen before storage at −80 °C. Tissue samples were crosslinked with 4% formaldehyde under vacuum on ice for 1 hour, and the reaction was quenched by adding 1 M glycine. Nuclei were isolated using NIB buffer (50mM HEPES pH7.4, 5mM MgCl2, 25mM NaCl, 5% sucrose, 30% glycerol, 0.25% Triton X-100, 0.1 % β-mercaptoethanol and 0.1% protease inhibitor), and chromatin was extracted in TE-SDS buffer. Chromatin was sheared by sonication for 30 s ON/30 s OFF cycles for a total of 25 cycles. For immunoprecipitation, the sonicated chromatin was incubated overnight at 4 °C with 2 ng of the following antibodies: anti-CENH3^70^, anti-H3K4me3 (Abcam, ab8580), anti-H3K9me2 (Abcam, ab1220), anti-H3K27me3 (Merck, 07-449). Control without antibody was performed simultaneously. The antibody–chromatin complexes were captured using rProtein A Sepharose® Fast Flow (Sigma, GE17-1279-01) or Protein G Sepharose® 4 Fast Flow(Sigma, GE17-0618-01). Bound chromatin was eluted, and de-crosslinked by Proteinase K treatment. DNA was purified by ethanol/sodium acetate precipitation and resuspended in nuclease-free water. The purified DNA was sent for library preparation and sequencing.

Sequencing reads were aligned to the reference genome using Bowtie2^71^ with the --very-sensitive-local option. ChIP and control alignments were compared using bamCompare^72^ with the parameters --binSize 50 --normalizeUsing RPKM -- operation log2. The resulting log_2_ ratio tracks were visualised using pyGenomeTracks^73^. Metaplots were generated using computeMatrix^72^ (scale-regions mode) with the parameters - -regionBodyLength 20000 --beforeRegionStartLength 10000 --afterRegionStartLength 10000 --binSize 50, and plotted using plotHeatmap.

### Probes design and oligo-FISH

After phased genome assembly and synteny analysis, a haplotype-specific translocation in *R. tenuis* reference plant was identified at the chromosome ends. In order to validate this data, we designed oligo probes specific to each translocated region and hybridized in the reference plant, in different accessions, plus *R. austrobrasiliensis*. Probes were designed for the following regions to each chromosome/haplotype: for chromosome 1 (Chr1_h1: 0-3100219bp/ Chr1_h2: 0-2085189bp) in a density of 3 probes per Kilobase Pair (Kb) (**Fig. 1c**), and for chromosome 2 (Chr2_h1: 133059273bp - end/ Chr2_h2: 136628596bp - end) in a density of 1 probe per Kb. After designing and setting the regions corresponding to each haplotype, this data was submitted to Daicel Arbor Biosciences (Michigan, USA) for synthesis and labelling of the probes. Probes for haplotype one were labelled in Atto633 (far red), while probes for haplotype two were labelled in Alexafluor488 (green). For the slides preparation, young roots (mitosis) and flower buds (meiosis and microgametogenesis), from plants maintained in pots at greenhouse facilities of Max-Planck Institute for plant Breeding Research, were collected and fixed in Carnoy solution (methanol:acetic acid; 3:1, v/v) for 2-24h at room temperature then immediately used or kept at -20 ºC until the moment of use. After fixation, the material was digested in enzymatic solution containing 2% cellulase Onozuka (Serva), 2% pectolyase Y-23 (Duchefa Biochemie), 2% cytohelicase (Sigma) and 10% pectinase (Sigma) in citrate-phosphate buffer (pH 4.5), washed in water and macerated in 60% acetic acid, post-fixed with ice cold fresh Carnoy solution and air dried. For removal of cytoplasm, the slides were washed in 60% acetic acid for at least 30 minutes, then air dried. For the pollen tube germination experiment, 30 mature flower buds were collected into 3 ml of sdH_2_O and vortexed to release pollen grains. From the bottom of the tube, 300 µl of the pollen suspension was collected and added to 3 ml of liquid medium for trinucleate pollen, following Tushabe and Rosbakh^74^. Pollen grains were incubated at 28 °C for 24 hours. After incubation, the medium containing germinating pollen grains was transferred into a conical tube and centrifuged at 5000 rpm for 3 minutes. After removing the supernatant, pollen nuclei were suspended in 60% AcAc, transferred onto a slide, post-fixed, and air-dried. For oligo-FISH, the procedures were similar to the ones published by Nascimento and Pedrosa-Harand^75^, briefly the hybridization mix was composed of 50% formamide (Sigma), 2×SSC (Saline Sodium Citrate) solution (pH 7.0), 10% dextran sulphate, 350 ng of probe labelled in Alexa Fluor-488 and 200 ng of the probe labelled in Atto-633 (a total of 15 µL per slide). After denaturation and hybridisation, the slides were passed through stringency washes in 2× and 0.1× SSC at 42 ºC, which corresponds to a final stringency of ∼76%. Slides were counterstained with 2 µg/mL DAPI in Vectashield H-100 antifade mounting medium (Vector Laboratories) and photographed on a Zeiss Microscope coupled with a Zeiss Axio Imager 2 camera system and software ZenBlue v3.2. Images were treated in brightness and contrast using Adobe Photoshop software.

### Immunocytochemistry

Immunocytochemistry was performed as described in Castellani et al.^18^, with some variations. Briefly, young flowers of *R. tenuis* and *R. austrobrasiliensis* were sampled and fixed in ice-cold 4% (w/v) paraformaldehyde (PFA) in phosphate buffered saline (PBS) solution (pH 7.5, 1.3 M NaCl, 70 mM Na2HPO4, 30 mM NaH2PO4) and 0.1% (v/v) Triton X-100 for 30 min in a vacuum. Flowers were dissected to select anthers of the appropriate size in order to have meiocytes in Prophase I. Anthers were opened from the tip in a drop of PBS with 0.1% (v/v) Triton X-100 and squeezed to release the meiocytes from the locules. The suspension of meiocytes was stirred to separate individual nuclei and remove excess debris. A coverslip was pressed onto the suspension to squash the meiocytes and later removed with liquid nitrogen. Slides were mounted with Vectashield containing 0.2 µg/ml DAPI and checked for Prophase I stages of interest. Selected slides were incubated with blocking buffer (3% (w/v) bovine serum albumin (BSA) in PBS + 0.1% (v/v) Triton X-100) for 1 hour at 37 °C to block and permeabilize the cells. The following antibodies were used: anti-AtASY1 raised in rabbit (inventory code PAK006)^23^, anti-AtMLH1 raised in rabbit (PAK017)^76^. The anti-RpZYP1 was raised in rat against the peptide KLTAERLVKDQASVKNDLEC (Gene ID: RP1G00482580/RP4G01479980/RP2G00810370/RP5G017 38420) and affinity-purified (Lifeprotein). The anti-RpREC8 was a combination of two antibodies raised in rabbit against the peptides CEEPYGEIQISKGPNM and CYNPDDSVERMRDDPG (Gene ID: RP1G00316120/R P2G00915110/RP4G01319620/RP5G016 38170) and affinity-purified (Eurogentec)^18^. The anti-RpHEI10 was a combination of four antibodies raised in rabbit and rat against the peptides CNRPNQSRARTNMFQL, CPVRQRNNKSMVSGGP, CIDIMDSRDMLRQGKREREEIW CDTDSAVNMGPPSGDTSNRR and (Gene ID: RP3G01271190/RP3G01008630/RP1G00269340/RP2G006 99130) and affinity-purified (Eurogentec and Lifeprotein)^18^. Primary antibodies were diluted in the blocking buffer to a final volume ratio of 1:200. Slides were incubated with primary antibodies overnight at 4 °C. The following day, slides were washed three times for 5 min each with PBS + 0.1% (v/v) Triton X-100. Samples were incubated with secondary antibodies for 2h at room temperature or 1h at 37 °C. Secondary antibodies were conjugated with STAR ORANGE (Abberior, STORANGE-1007) or Alexa Fluor 488 (Thermofisher Goat anti-Rabbit IgG (H+L), Superclonal™ Recombinant Secondary Antibody, A27034) diluted 1:250 in blocking buffer. Slides were washed again three times for 5 min with PBS + 0.1% (v/v) Triton X-100 and allowed to dry. Samples were then prepared with 10 µl of mounting solution (Vectashield + 0.2 µg/ml DAPI). Specimens were covered with a coverslip and sealed with nail polish for storage. Images were taken with a Zeiss Axio Imager Z2 with Apotome system for optical sectioning. Images were deconvolved and processed with Zen 3.2 software and Adobe Photoshop.

### Meiotic gene analysis

Meiotic genes with a role specifically in meiotic recombination were selected from the literature based on plants and other eukaryotes. Sequences were selected from *Arabidopsis* or closer relatives in the case of poorly conserved genes (e.g. rice, maize, barley). Protein sequences were used as a query for tblastn with our in-house genomes, annotations and expression datasets. The number of hits on the same query sequence was used as a starting measure of copy number. Results were then manually curated for validation (**Supplementary Fig. 11**). Expression evidence for meiotic genes was based on the analysis of the inflorescence transcriptome obtained by RNA-seq data.

To further evaluate *SHOC1* transcriptional activity, total RNA from 100mg of flower buds was isolated using the Spectrum Plant Total RNA Kit (Sigma 000393619), followed by cDNA synthesis with the SuperScript IV Reverse Transcriptase (ThermoFisher 18090050). The obtained cDNA was used as a template for PCR with Phusion HF DNA Polymerase (NEB M0530), successfully amplifying the predicted coding sequence of *SHOC1*.

### Pollen Nuclei Isolation and Sequencing

The protocol was adapted from Castellani, et al.^18^. For collecting the pollen, mature flowers of *Rhynchospora tenuis* were collected into 5ml tubes containing Woody Pollen Buffer (scWPB: 200mM Tris-HCl; 4mM MgCl_2_; 2mM EDTA; 86mM NaCl; 10mM Na_2_S_2_O_5_; 250mM Sucrose; 0,5mM Spermine; 0,5mM Spermidine; 1% PVP-10), and vortex for 30s at maximum speed. The obtained solution was filtered through a 50µm CellTrics cell strainer into a 5ml tube. Samples were centrifuged at 5000g for 5 minutes, the supernatant was removed, and the pollen pellet was frozen in liquid nitrogen. Pollen viability was assessed by Alexander staining (Morphisto 13441-00250). Pollen samples were stored at -70ºC until further use.

For extracting the pollen nuclei, frozen pollen samples were thawed on ice for 5 minutes and resuspended in scWPB. The solution was filtered through a 5µm CellTrics cell strainer, retaining the pollen. Pollen grains inside the cell strainer were crushed with a plastic rod. Pollen nuclei were filtered from the macerated pollen by adding scWPB with some additives to preserve RNA (scWPC; 5mM DTT; 1%BSA; 0.2 U/µL Protector RNase Inhibitor). Pollen nuclei solution was stained with DAPI (1µg/µl) and used for Fluorescence-activated cell sorting (FACS) to enrich the nuclei population. BD FACSAriaIII Fusion Flow Cytometer with a 70µm nozzle and 483 kPa sheath pressure was employed for this purpose. Nuclei were dispensed into a 96-well plate containing Collection Buffer (1xPBS, 1%BSA, and 0.2 U/µL Protector RNase Inhibitor). The quality and number of nuclei were evaluated with the LUNA-FX7 Automated Cell Counter. Nuclei solutions were used for library preparation using Chromium Next GEM Single Cell ATAC or Chromium Next GEM Single Cell 5’ Kits following the manufacturer’s instructions. Libraries sequenced at BGI Genomics following Chromium 10X Kit recommendations.

### Single-cell analysis

Single-cell RNA data were analysed with a similar pipeline reported in the previous study in *R. breviuscula*^18^. A full description of the pipeline was updated in the GitHub page: https://github.com/Raina-M/detectCO_by_scRNAseq.

### Subgenome-aware phasing of R. tenuis haplotypes

We used SubPhaser^21^ (default parameters) to phase and partition the haplotypes of *R. tenuis* by assigning chromosomes to subgenomes based on differential repetitive *k*-mers. These were assumed to have expanded during the period of independent evolution after the loss of recombination. A subgenome is considered to be well phased when it displays distinct patterns of both differential *k*-mers and homoeologous chromosomes, confirming the presence of subgenome-specific features.

### Estimating divergence times between haplotypes

Divergence times between haplotypes were estimated using the method outlined in Guo et al.^77^. In brief, all SNPs, derived from SyRI (v1.5.3)^64^ analysis explained above, between two haplotypes were used to estimate divergence times according to the formula *g=d/2μ*, where *g* is the number of generations, *μ* is the assumed mutation rate of 6.13×10^-9 78^, and *d* is the number of SNPs per bp (*d*=*total number of SNPs between two haplotypes/chromosome size*). We assumed a generation time of one per year.

LTR insertion times were calculated by Subphaser as follows: LTR-TRs were de novo detected using LTRharvest (v.1.6.1)^79^ and LTRfinder (v.1.07)^80^. To reduce false positives, TEsorter (v.1.3.0)^81^ was used to reconstruct the classification of LTR-RTs and further refine this classification. The subgenome-specific *k*-mer sequences were mapped to the LTR-RT sequences using a substring match procedure to identify the subgenome-specific LTR-RTs using Fisher’s exact test. Two LTRs of each subgenome-specific LTR-RT were retrieved and the nucleotide divergence was estimated using the Jukes–Cantor 1969 model. The insertion time (*T*) was calculated using the equation *T* = *K*/2*r*, where *r* = 6.13×10^-9^ substitutions per year, and *K* represents the divergence of the LTRs from the LTR-RT.

*Ks* between two haplotypes was computed based on the CDS sequences annotated by Helixer by CoGe (https://ghibli.bti.cornell.edu/coge/). Genomes and gene annotations were uploaded to CoGe, and the option to calculate *Ks* was selected during the SynMap run. All *Ks* of coding sequences that overlapped with transposable elements annotation was removed by ‘bedtools subtract -A’.

### Phylogenetic tree construction

The phylogenetic tree was built based on single copy orthologs (SCOs) among *R. breviuscula, R. austrobrasiliensis*, and all haplotypes of *R. tenuis*. SCOs were extracted from the output of OrthoFinder in GENESPACE. All SCO groups were aligned by MAFFT (v7.475)^82^ with parameter ‘--auto –maxiterate 10’. All alignments were concatenated with a custom python script available in our project GitHub page (phylogeny_based_on_SCOs/concatenate_aln.py). The tree was constructed based on the catenated alignments with RAxML-NG (v1.2.2)^83^, then visualised in FigTree (v1.4.4) (https://github.com/rambaut/figtree/).

### F1 genotyping and analysis

F1 individuals were obtained from controlled self-crossed heterozygous mother plants. Genomic DNA from seedlings was extracted and sequenced on an Illumina NovaSeq platform. Reads were mapped to haplotype 1 assembly, and SNP genotypes were called using bcftools (v1.9). Progeny haplotypes were reconstructed and compared to maternal alleles to assess inheritance patterns and identify crossover events. F1 genotyping method was the same as Castellani et al.^18^.

### Embryo and endosperm analysis

For extracting the nuclei from embryo and endosperm, frozen seeds were macerated with a plastic rod and resuspended in Seed Nuclei Isolation Buffer (100mM Tris, 5.2 mM MgCl2, 85.55 mM NaCl, 0.1% Triton, 0.2 U/µL Protector RNase Inhibitor, pH 7.5). The obtained solution was filtered through a 20µm CellTrics cell strainer into a fresh 2ml tube and stained with DAPI (1 µg/µl). BD FACSAriaIII Fusion Flow Cytometer with a 70µm nozzle was employed for FACS to enrich the 2c (embryo) and 3c (endosperm) nuclei population. The separate 2c and 3c nuclei populations were dispensed into a 96-well plate containing Collection Buffer (1xPBS, 1%BSA, and 0,2 U/µL Protector RNase Inhibitor). The quality and number of nuclei were evaluated with the LUNA-FX7 Automated Cell Counter. Nuclei solutions were used for library preparation using Chromium Next GEM Single Cell ATAC following the manufacturer’s instructions. Libraries sequenced at BGI Genomics using the Chromium 10X Kit recommendations.

### Female gametophyte sectioning and imaging

Developing ovaries and whole inflorescences were fixed according to Rocha et al.^30^. Dehydration and infiltration into Araldite 502/Embed 812 resin were performed in an EMS LYNX II automated tissue processor (EMS, Hatfield, PA, USA). Following resin polymerisation, serial semithin sections (1 µm) were produced from ovules at different stages of development, stained with 1% aqueous toluidine blue supplemented with 1% sodium tetraborate and imaged with a Zeiss Axio Imager M2. Images were stacked and aligned in FIJI using the StackReg plugin (BIG-EPFL). MorphoGrahX was used for 3D rendering of the aligned images and volumetric visualization of the embryo sac tissue^84^. 3D segmentation of the embryo sac was performed using ITK watershed (https://www.itk.org) in MorphoGraphX, taking PlantSeg^85^ predictions from “generic confocal 3DUNET model” as input images for 3D cell segmentation. Immunodetection of ß-1,3-glucan to identify pollen tube cell walls was done as described in Freh et al.^86^, with the exception that BSA was omitted from the buffer during all steps. Goat anti-mouse Alexa Fluor 647 (abcam ab150119, Cambridge, UK) was used as an alternative secondary antibody. Imaging was done on a Leica SP8.

For Feulgen staining of siliques, the pericarp was removed from the mature and fertile siliques; for the rest of the siliques, the pericarp was not removed, and the material was fixed in ethanol:acetic acid (3:1) overnight. The samples were washed three times with distilled water for 15 min each wash and then incubated in 5 N HCl for 1 h, followed by another series of three washes with water, as before. The siliques were then incubated in Schiff’s reagent for 3–4 h, after which they were washed three times with cold distilled water (4°C). Next, the siliques were incubated in ethanol series 10 min each, 10%, 30%, 50%, 70%, 95% and then washed several times with 99.5% ethanol until the ethanol came out colorless. The samples were incubated in 99.5% Ethanol:LR White series 15 minutes each solution 3:1, 2:1. The samples were then incubated for 1 hour in ethanol:LR White resin (1:1), followed by an overnight incubation in LR White resin. Afterwards, the seeds were mounted on microscope slides in LR White resin and polymerized for 24h at 60°C. The samples were imaged under Multiphoton microscopy, Leica Stellaris 8 and Leica SP8 Falcon Dive with emission around 580 nm and excitation between 580 nm - 700 nm.

## Data availability

Genome assemblies and raw sequencing data have been deposited in the European Nucleotide Archive under accession PRJEB98016. The reference genomes, sequencing data, annotations and all tracks presented in this work are made available for download at EDMOND, the Open Research Data Repository of the Max Planck Society. All other data are available from the corresponding author upon request.

## Contributions

A.M. conceived the research and supervised the project. M.Z., P.G.H. and A.M. performed the genome assembly, scaffolding and genomics. M.Z., J.C., H.S., M.C., and S.S. designed and performed the single-cell experiments and processed the data. K.S. assisted the single-cell analysis. M.C. performed the immunostaining experiments. L.D. and M.C. performed the functional screening of meiotic genes. T.N. and M.Majka performed FISH experiments. M.Majka performed pollen-FISH genotyping experiments. G.T. and M.Z. performed the ChIP–seq analysis. M.Marek and S.S. performed the flow cytometry measurements and nuclei sorting. N.S., M.Marek and B.H. performed all nuclear sequencing libraries. M.Z., L.A.R., A.L.L.V. and A.M. performed the centromere and repeat characterisation. A.L.L.V. collected and provided the plant material. U.P. and M.Majka performed the plant crosses. S.S. performed the pollen viability assay. M.Z., S.D. and A.M. performed the dating analysis. M.Z., T.L. and K.F.X.M. performed the functional annotation. G.T., U.N., A.V., T.T. and D.F. performed and analysed the female gametophyte and embryo sac development microscopy data. M.Z., M.C., S.S. and A.M. wrote the first manuscript draft with input from all authors. All authors approved the final version of the manuscript.

## Acknowledgements

This study was funded by the Max Planck Society (core funding to A.M.), the German Research Foundation (MA 9363/2-1 and MA 9363/3-1), and the European Union (European Research Council Starting Grant, HoloRECOMB, grant no. 101114879 to A.M.). The DFG also funded this work under Germany’s Excellence Strategy—EXC 493 2048/1–390686111 (to K.S. and A.M.). M.Z. is financially supported by the DFG (grant no. MA 9363/2-1). M.Majka is supported by the Alexander von Humboldt Foundation (Humboldt Research Fellowship for Postdoctoral Researchers). G.T. is a recipient of the predoctoral fellowship HORIZON-MSCA-2021-COFUND-01, rePLANT GA 101081581, funded by the European Union and co-funded by the Max Planck Society. The views and opinions expressed are those of the author(s) only and do not necessarily reflect those of the European Union or the European Research Executive Agency (REA). Neither the European Union nor the granting authority can be held responsible for them.

## Extended Data Figures

**Extended Data Fig. 1.**
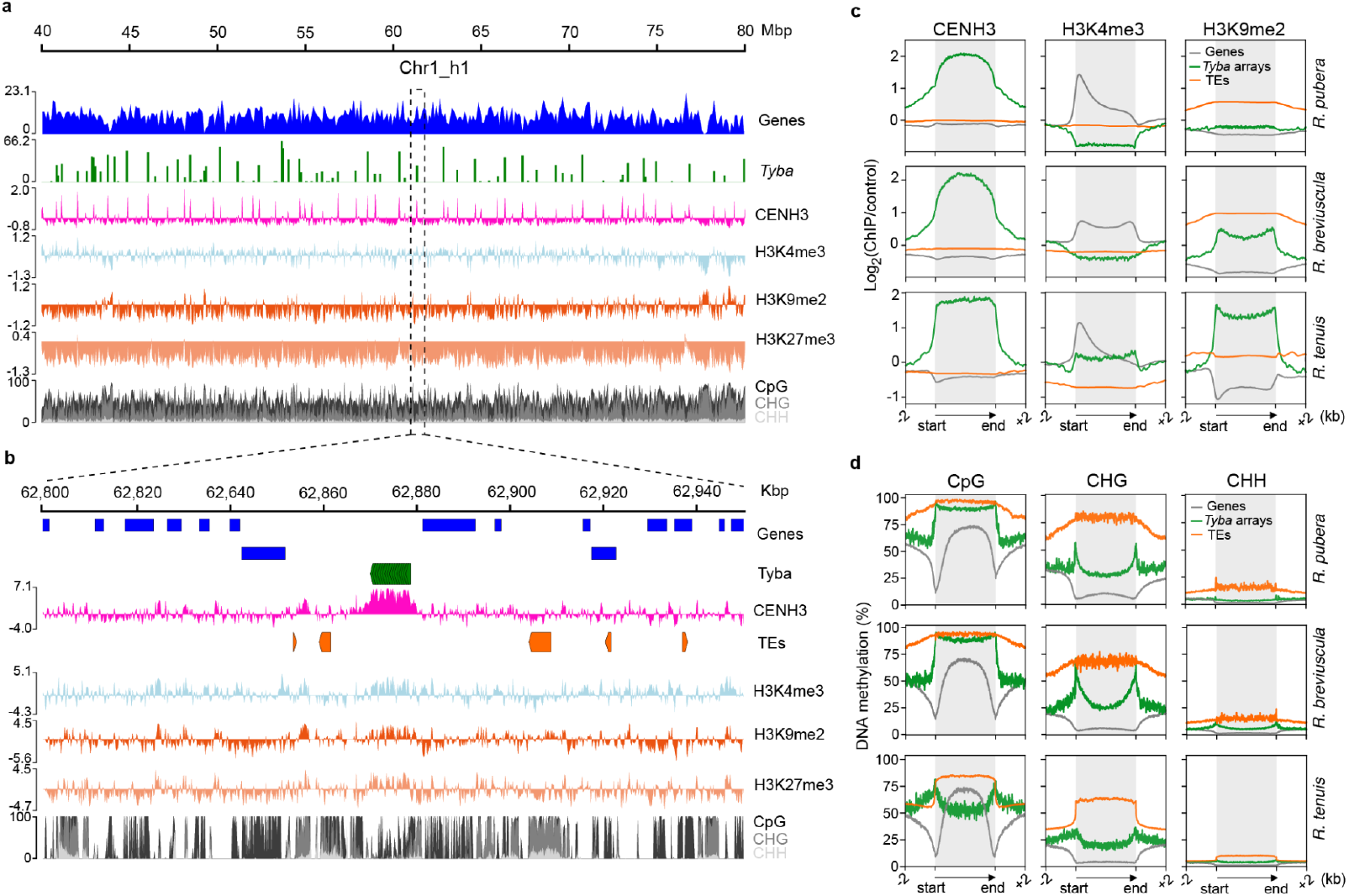
Divergent epigenetic landscape of holocentromeres in *Rhynchospora tenuis*. (**a**) Zoomed-in view of *R. tenuis* Chr1_h1 showing a 40-Mb region with multiple CENH3 domains that are closely correlated with *Tyba* repeat distribution. Gene and *Tyba* densities were calculated over 100-kb windows. (**b**) A detailed view of a representative ∼150 kb holocentromeric region on *R. tenuis* Chr1_h1, illustrating the precise correlation between CENH3 binding, *Tyba* arrays and an ambiguous chromatin state, with low CpG DNA methylation and accumulation of both repressive and active histone marks. Values in **a** and **b** represent the log2 ratio of ChIP-seq signal to input control. (**c**) Enrichment of CENH3, H3K4me3, and H3K9me2 from the start and end of different types of sequences: genes (gray line), TEs (orange) and *Tyba* repeats (green) for *R. pubera, R. breviuscula* and *R. tenuis*. ChIP-seq signals are shown as log2 (normalized RPKM ChIP/input). (**d**) Enrichment of DNA methylation in the CpG, CHG, and CHH contexts for the same sequence types as shown in (**c**) for *R. pubera, R. breviuscula* and *R. tenuis*. Gray boxes in (**c–d**) highlight the modification enrichment over the body of each sequence type.

**Extended Data Fig. 2.**
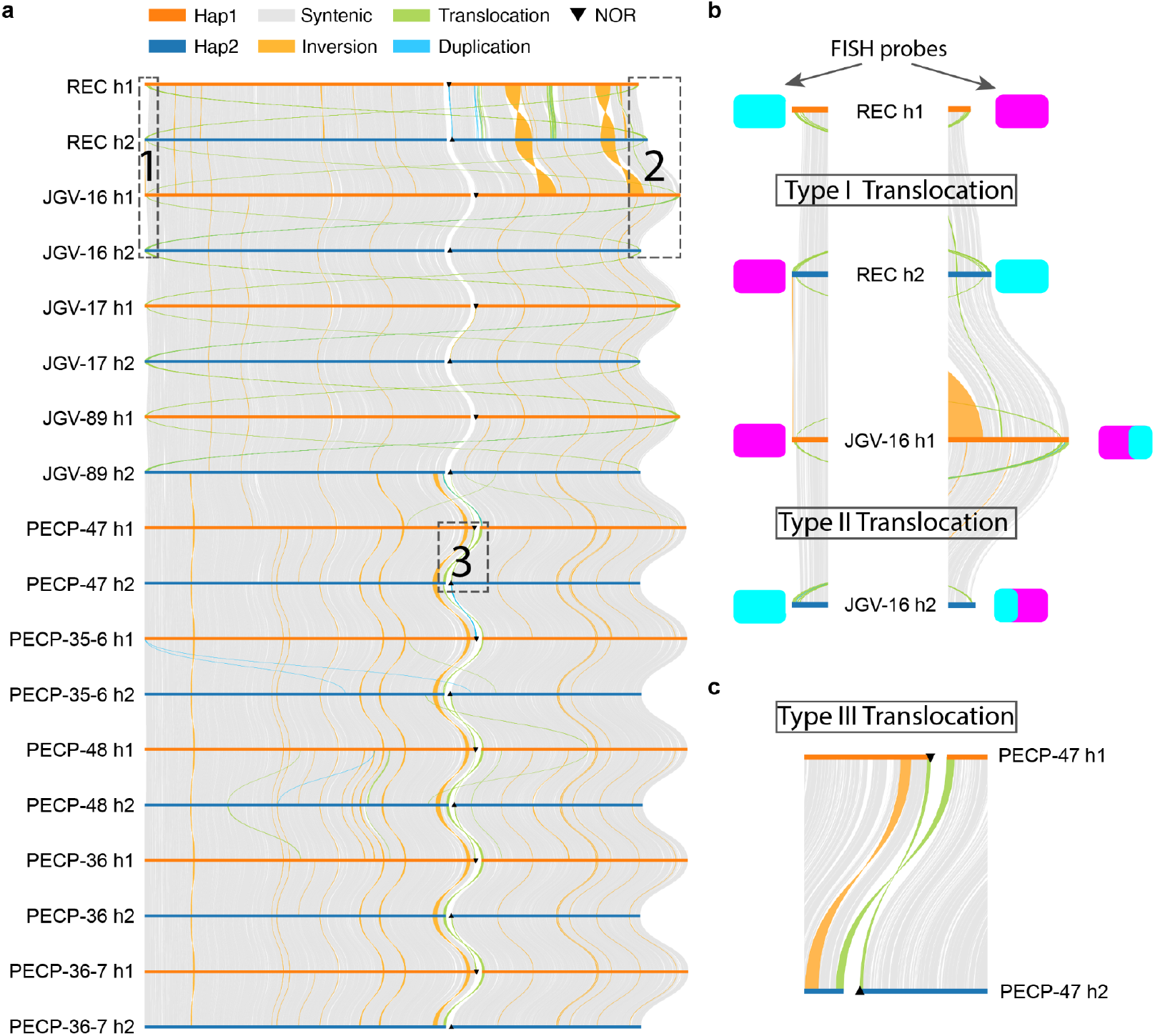
Colinearity across the two haplotypes of all *R. tenuis* accessions by SyRI. (**a**) Colinearity based on genomic sequences of all haplotypes of nice *R. tenuis* accessions (**b**) Zoom-in view of the regions 1 (left column) and 2 (right column) labeled in **a**. The translocation FISH probes designed for REC were also shown here, implying the Type I translocation. JGV-16 Chr 2 end (region 2) has a small translocation compared to REC Chr1_h2, so a mixed magenta and cyan probe signal was observed, indicating a different type of reciprocal translocation (Type II translocation). (**b**) Zoom-in view of the regions 3 labeled in **a**. (**c**) This displays the Type III translocation that found in PECP accessions, where the first reciprocal translocation is between Chr1_h1 end and Chr2_h2 start, and the second reciprocal translocation is between Chr1_h2 end and Chr2_h1 start.

**Extended Data Fig. 3.**
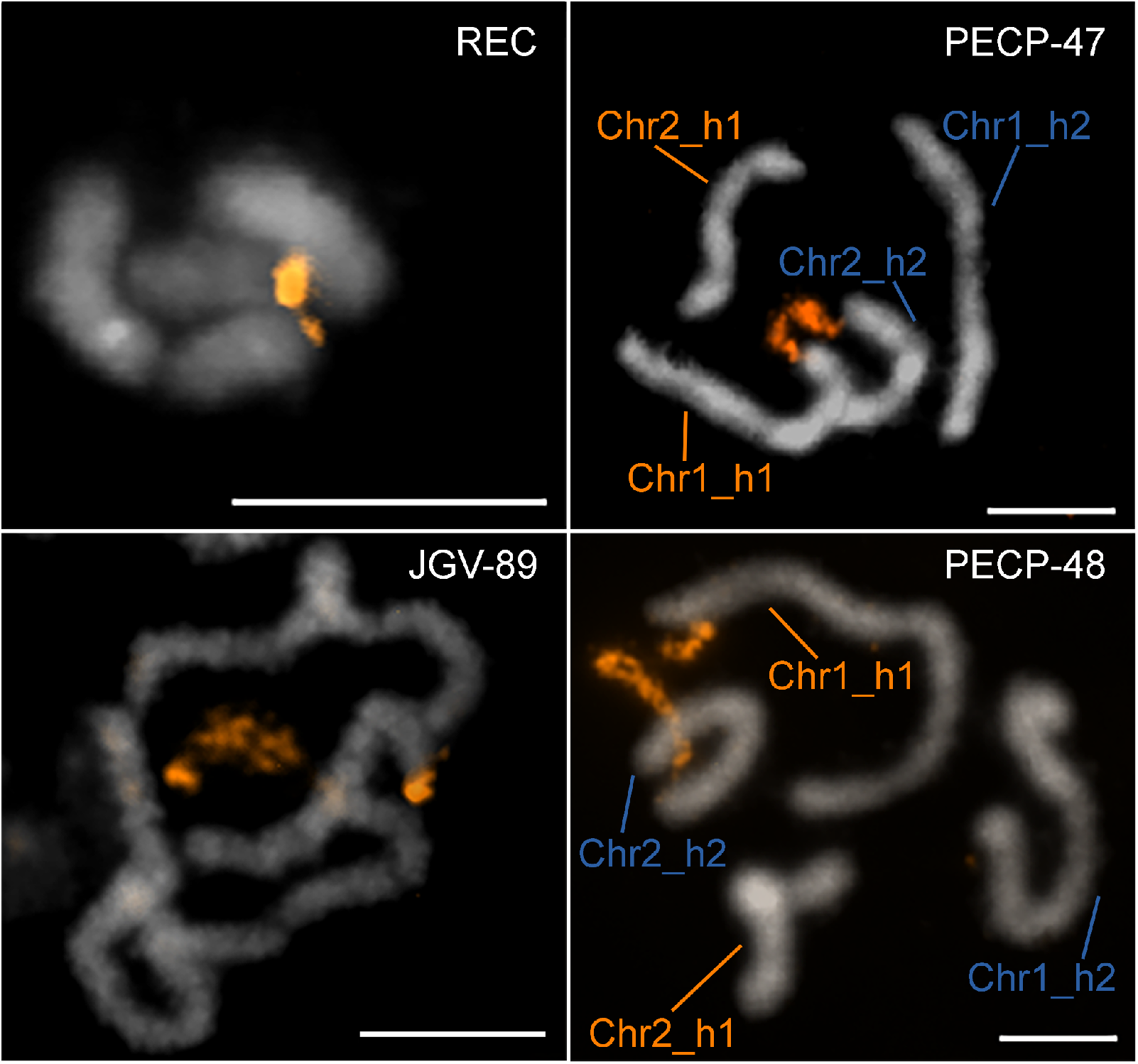
Distribution of 45S rDNA and identification of a reciprocal translocation in *R. tenuis* genotypes. FISH with a 45S rDNA probe (orange) on mitotic chromosomes of *R. tenuis* genotypes REC, JGV-89, PECP-47 and PECP-48. Genotypes PECP-47 and PECP-48 exhibit a reciprocal translocation involving 45S rDNA regions. The translocation occurs between chromosomes Chr1_h1 and Chr2_h2 and is absent in genotypes REC and JGV-89. Scale bars correspond to 5 µm.

**Extended Data Fig. 4.**
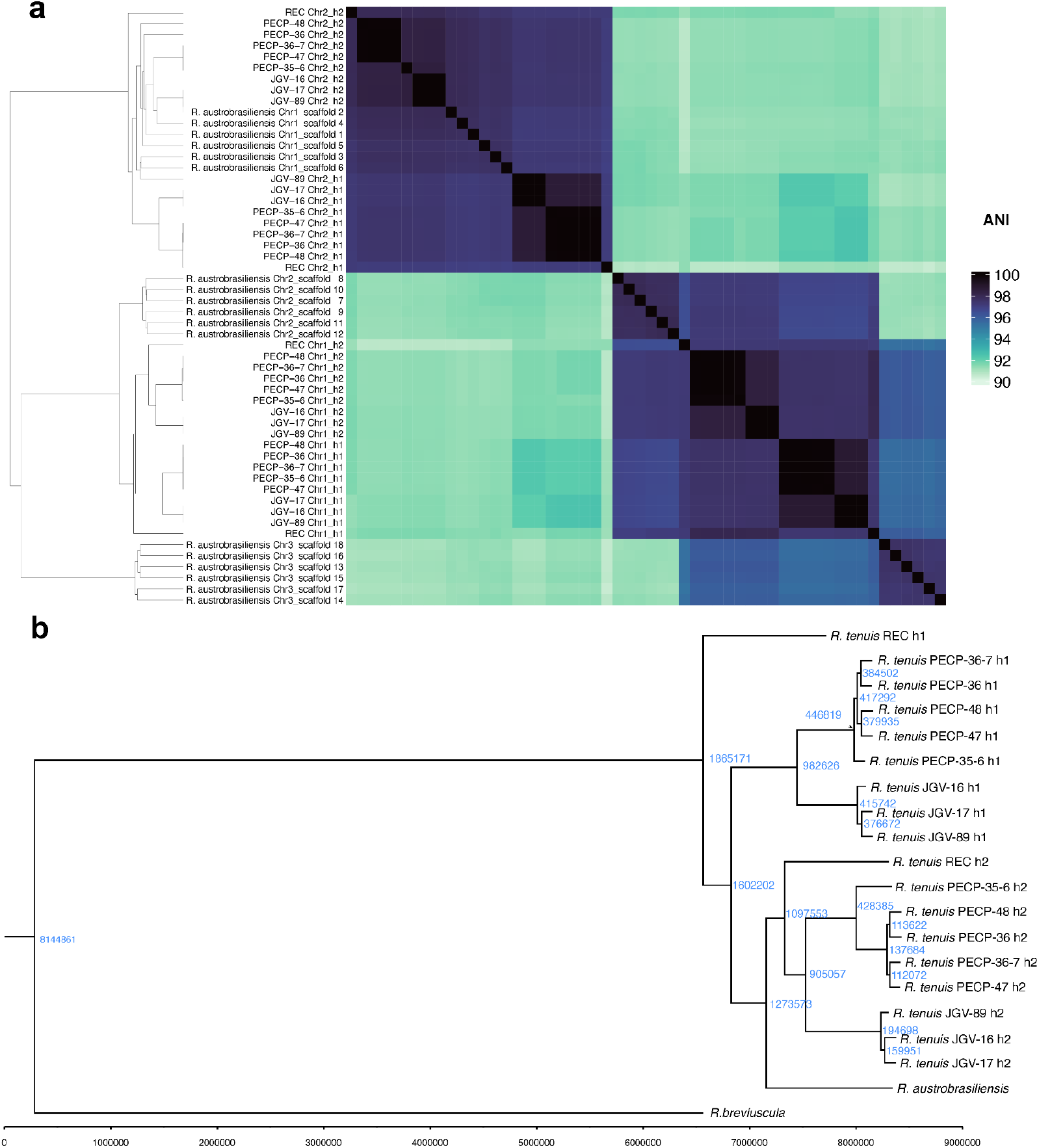
*K*-mer clustering and adjacency matrix of all 36 *R. tenuis* chromosomes from all accessions and 18 *R. austrobrasiliensis* pseudo-chromosomes. (**a**) *K*-mer size is 31 and the color bar on the right side indicates the similarity level measured by jaccard index. ANI: average nucleotide identity. (**b**) Phylogenetic tree rooted by *R. breviuscula* based on divergence of single-copy ortholog genes. All nodes were labeled with split time (blue number) of the corresponding branches. Time was calculated by *substitution per site* / (2 x *mutation rate per site per generation*). Mutation rate is 6.13e-9. Please note that this tree was constructed with only a pseudo-haplotype of *R. austrobrasiliensis* because the six haplotypes were not fully resolved. Here we used scaffold 1, 7, and 13.

**Extended Data Fig. 5.**
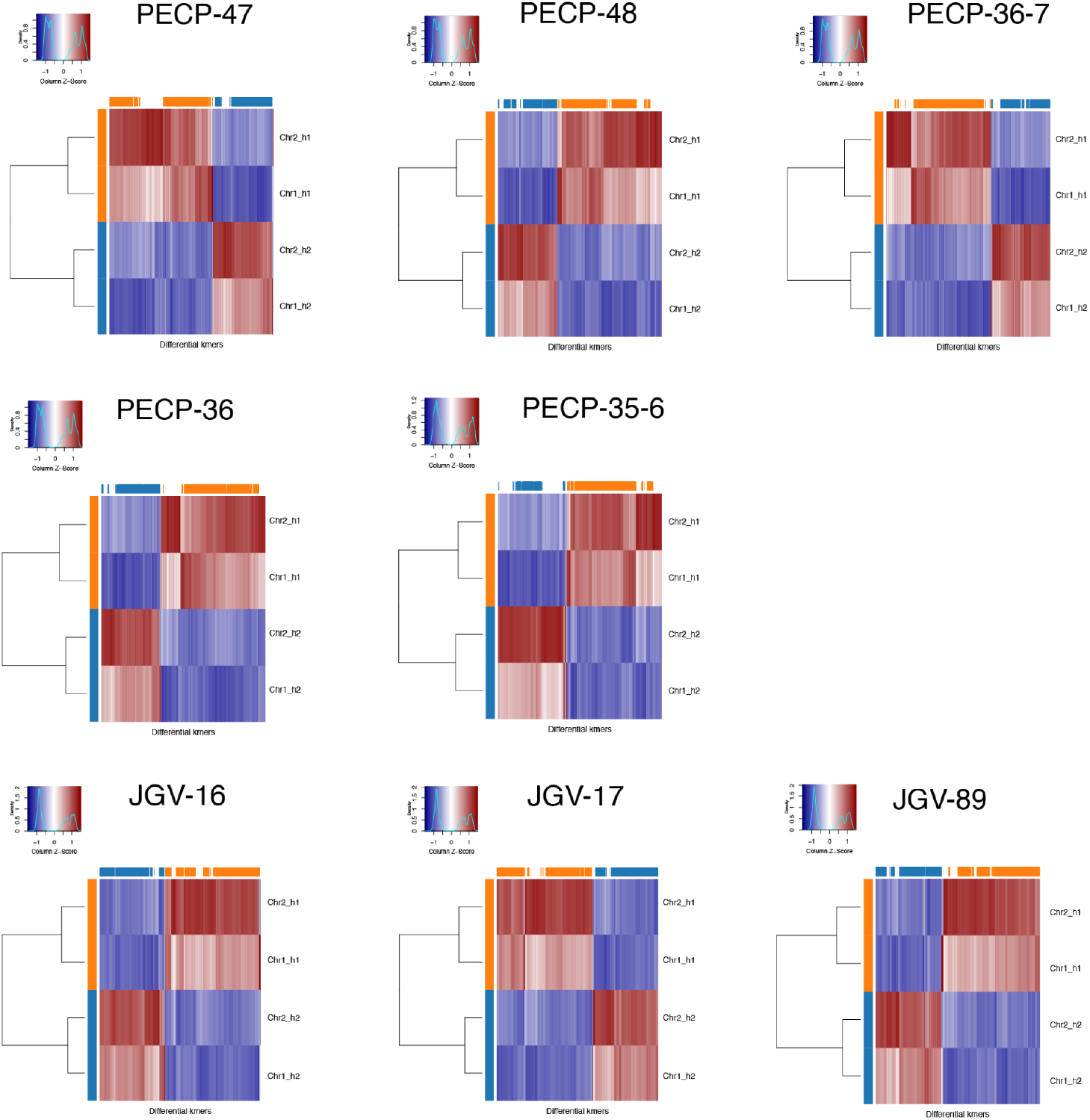
SubPhaser on phased haplotypes of different *R. tenuis* accessions. Unsupervised hierarchical clustering (the horizontal colour bar at the top of the axis indicates to which subgenome the *k*-mer is specific; the vertical colour bar on the left of the axis indicates the subgenome to which the chromosome is assigned). The heatmap indicates the *Z*-scale relative abundance of *k*-mers. The larger the *Z* score is, the greater the relative abundance of a *k*-mer).

**Extended Data Fig. 6.**
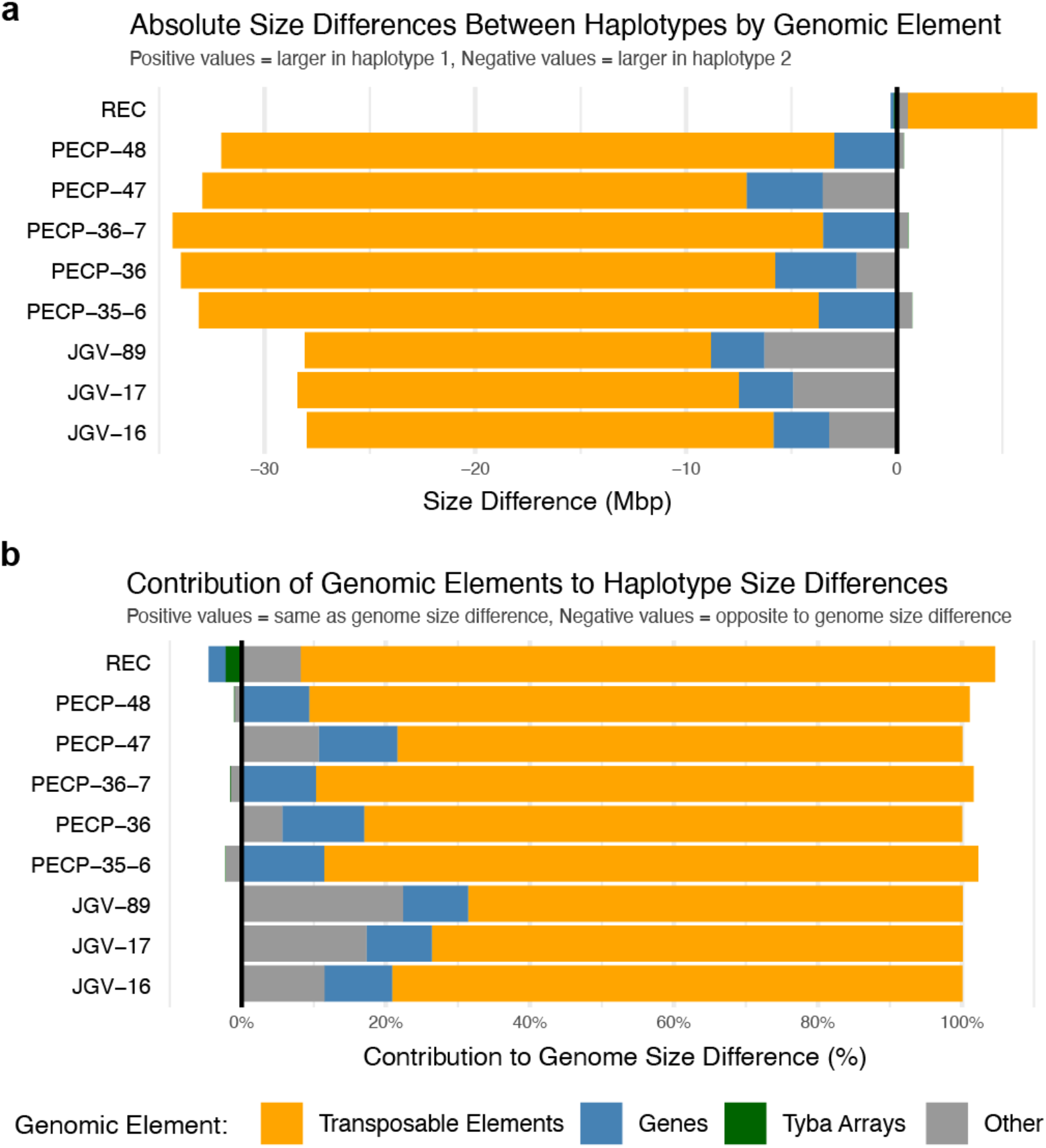
Genomic elements that contribute to the haplotype differences of all *R. tenuis* accessions. (**a**) Absolute size differences of different genomic elements between the two haplotypes of all *R. tenuis* accessions. A positive value means the total size of the element is larger in haplotype 1 and a negative value means a larger total size in haplotype 2. (**b**) Contribution of the genomic elements to the percentage of genome size difference of two haplotypes of *R. tenuis*. A positive value means this element difference is the same direction of genome size difference. For instance, haplotype 1 genome size of REC is smaller than haplotype 2. Transposable elements (TEs) in REC haplotype 1 is also shorter than haplotype 2, so the contribution of TEs has a positive value; while gene total length is longer in haplotype 1 so the contribution is negative.

**Extended Data Fig. 7.**
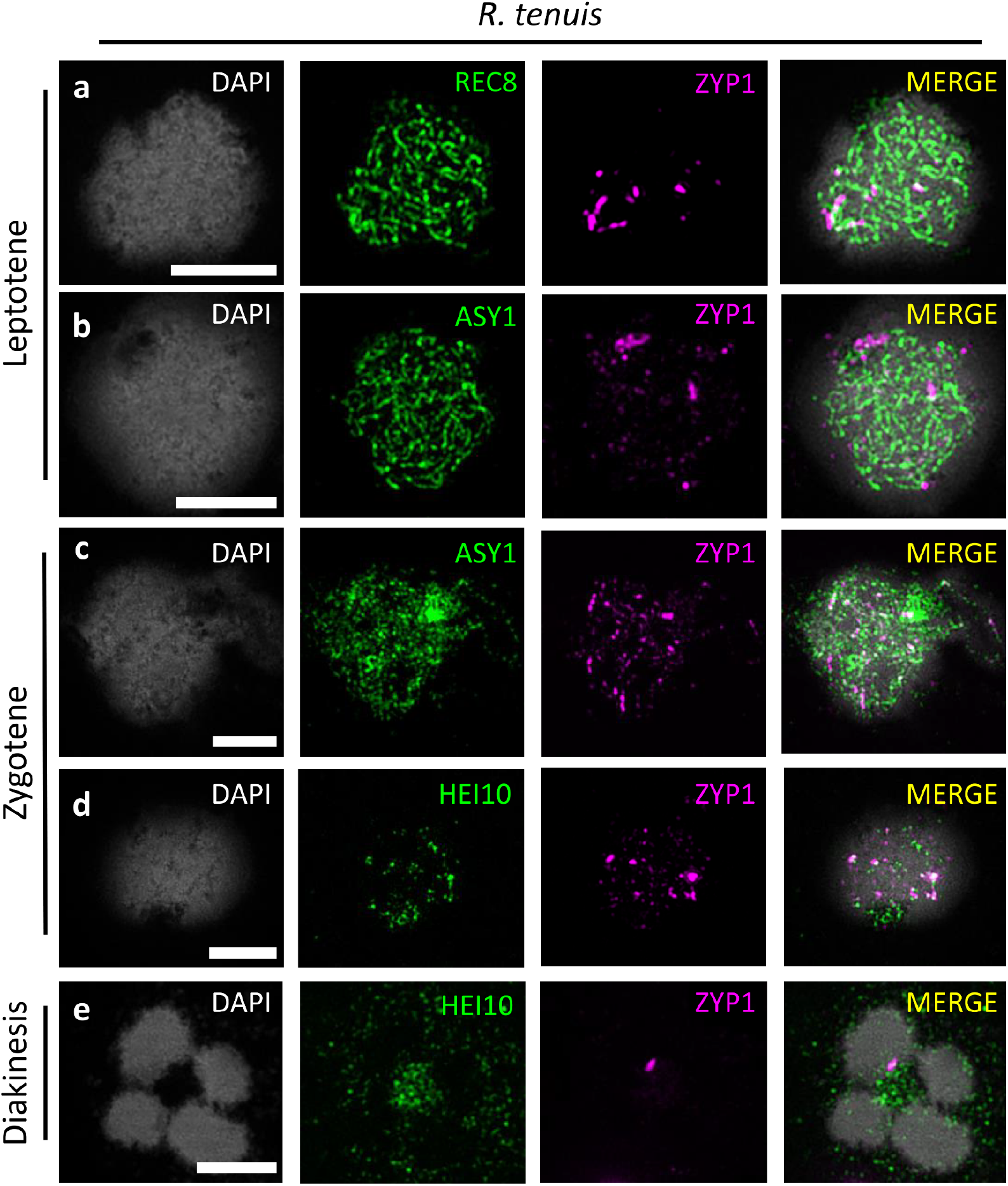
Additional channels and annotations of main figure, showing the meiotic atlas of *R. tenuis* and major recombination proteins. Early recombination proteins ASY1 and REC8 display regular behaviour, localising as linear signals (a-b). At zygotene, ASY1 is unloaded but ZYP1 shows only fragmented association with the axis. ZYP1 is able to attract unspecific signals of HEI10 (c, d). At diakinesis, HEI10 doesn’t localise as foci and four univalents are observed. ZYP1 forms polycomplexes. Scale bars correspond to 5 µm.

**Extended Data Fig. 8.**
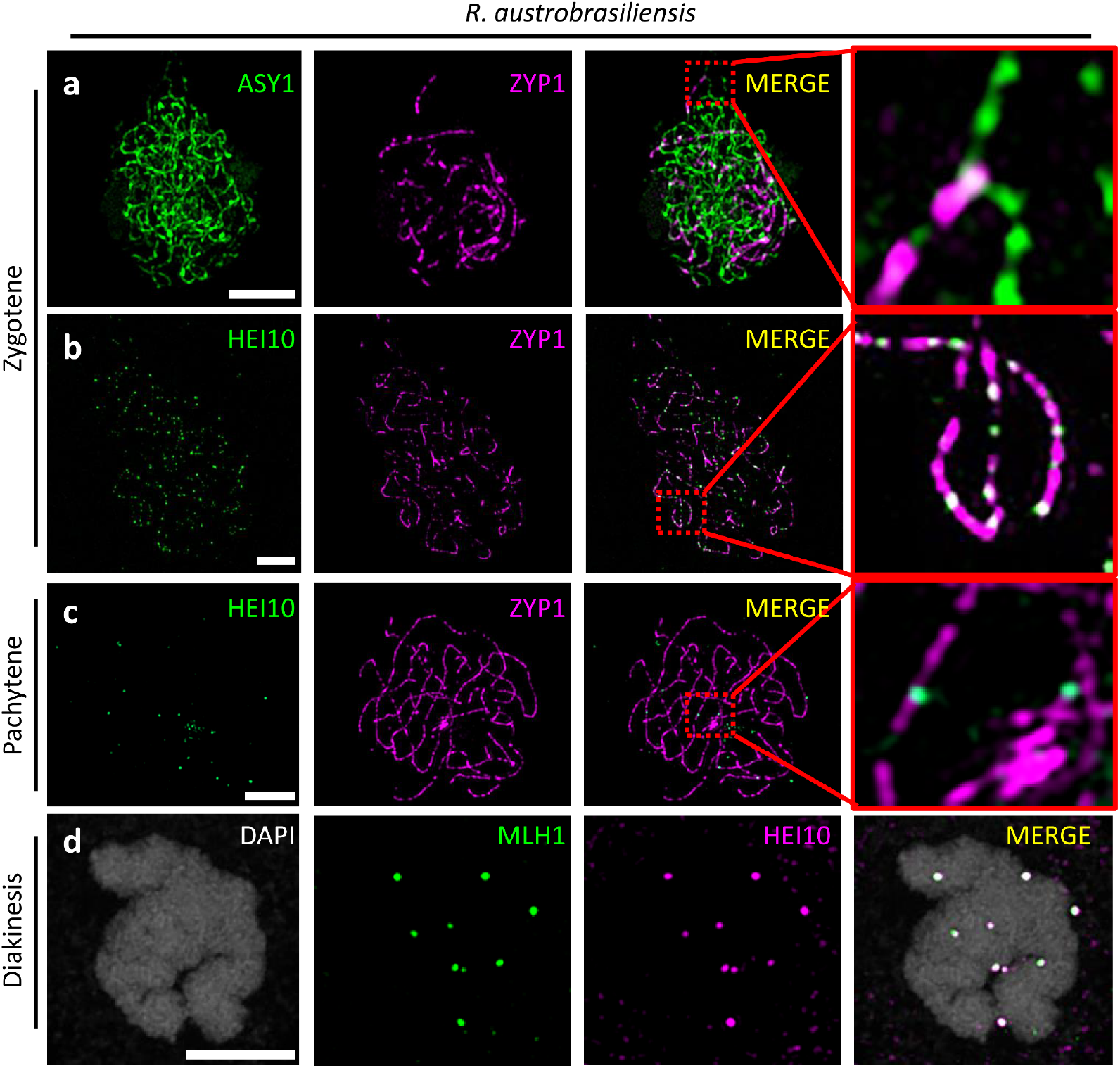
Additional channels and annotations of the main figure of *R. austrobrasiliensis* prophase I. At zygotene, ZYP1 is loaded on paired chromosomes while ASY1 is unloaded (**a**). Simultaneously, HEI10 is loaded as multiple foci along the signal of ZYP1 (**b**). At pachytene, ZYP1 forms a continuous linear signal along the whole chromosome length and HEI10 signal condenses into few high intensity foci (**c**). At diakinesis, HEI10 forms foci that colocalize with MLH1 and mark class I crossovers. Scale bars correspond to 5 µm.

**Extended Data Fig. 9.**
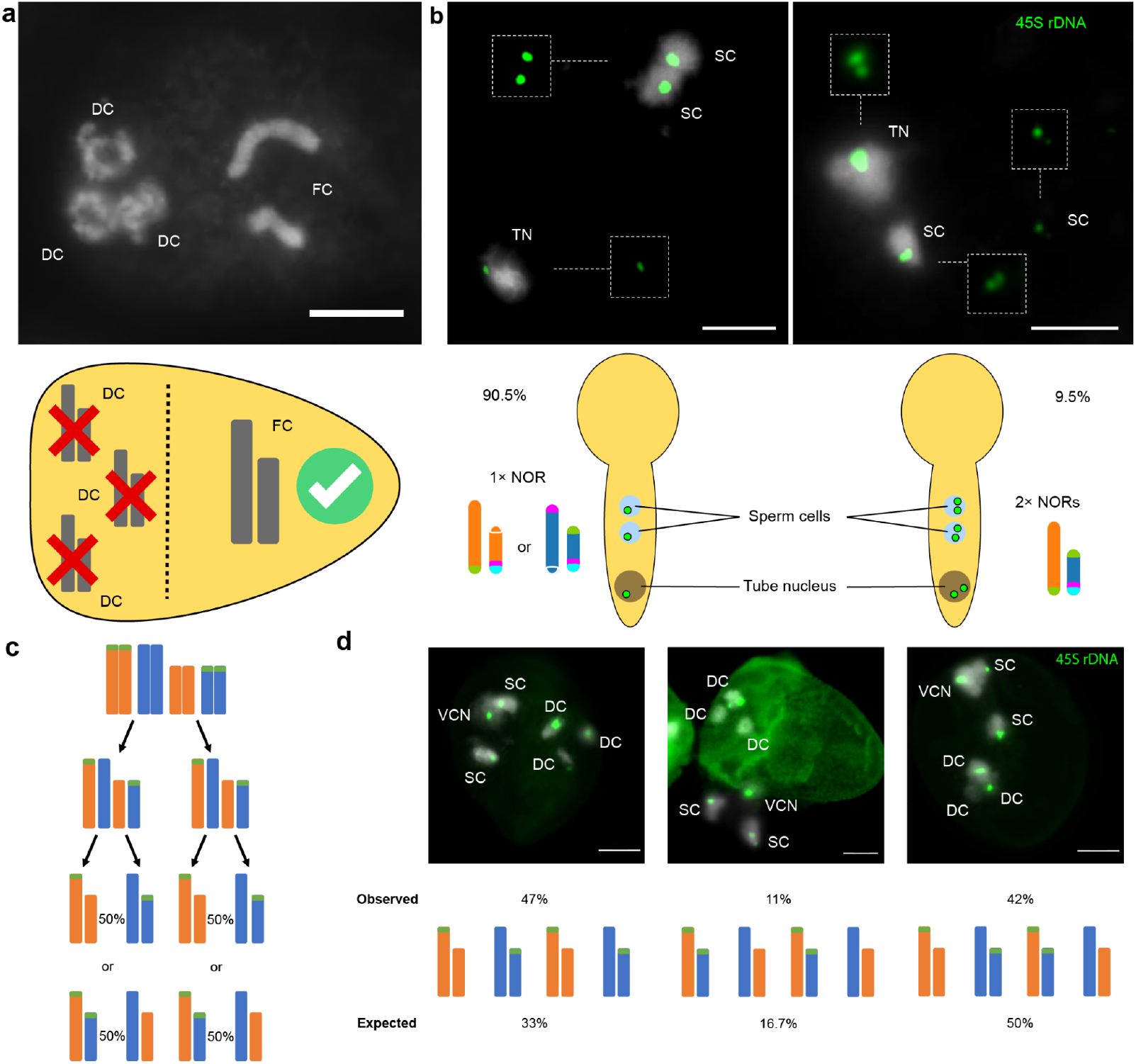
PECP-48 segregation distortion of NOR-bearing chromosomes during asymmetric male meiosis. (**a**) Male meiosis resulting in the formation of a single functional microspore (functional cell, FC) and three degenerating cells (DC), characteristic of monad-type microsporogenesis. (**b**) FISH detection of 45S rDNA loci (green) in sperm cells and tube nucleus of pollen tubes. Pollen grains exhibit either one (90.5%, left) or two (9.5%, right) 45S rDNA foci. The schematic representation below illustrates the relative frequencies of each class (90.5% and 9.5%) and possible chromosome haplotypes. (**c**) Predicted segregation outcomes and expected gamete genotype frequencies following achiasmatic inverted meiosis. (**d**) Expected versus observed (n = 100) frequencies of gamete genotypes based on FISH analysis of mature pollen grains hybridised with a 45S rDNA probe (green), illustrating segregation distortion favoring the larger, TE-rich haplotype 1. DC = degenerating cell; FC = functional cell; SC = sperm cell; TN = tube nucleus; VCN = vegetative cell nucleus. Scale bars correspond to 5 μm.

**Extended Data Fig. 10.**
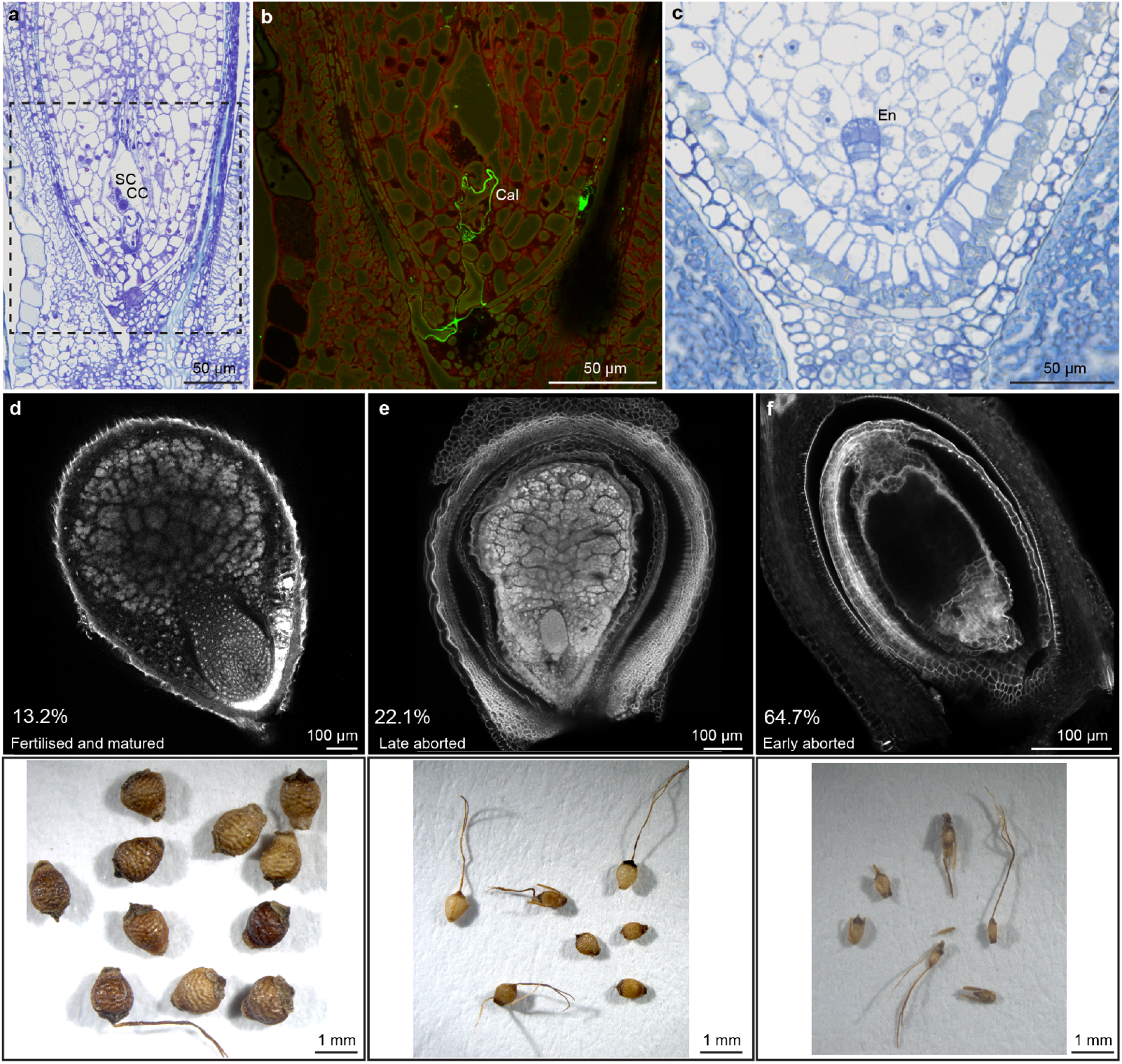
Embryology of *R. tenuis* (PECP-48). (**a**) Pollen tube entering the mature embryo sac to deliver the sperm cells. SC = sperm cell, CC = central cell. (**b**) The presence of the pollen tube in the ovule tissue is confirmed by the detection of callose (Cal) in green in the pollen tube cell wall (indirect immunofluorescent labeling of ß-1,3-glucan; see **Supplementary Figure 9**). (**c**) Abnormal development of seed with an embryo (En) but no visible endosperm formation. (**d**) Viable mature, (**e**) late aborted and (**f**) early aborted seeds shown by Feulgen staining (top) and respective seed morphology (bottom).

