## Supplementary Material for "Sex without crossovers mimics clonal reproduction in the holocentric plant *Rhynchospora tenuis*"

### Supplementary Tables

**Supplementary Table 1.** Genome assembly statistics of all *R. tenuis* accessions and *R. austrobrasiliensis*

**Supplementary Table 2:** List of meiotic genes searched in *R. tenuis* genotypes, their location and comparison with *R. breviuscula* and *R. austrobrasiliensis.*

**Supplementary Table 3.** Cell number of snRNA- and snATAC-seq of different *R. tenuis* accessions.

### Supplementary Figures

**Supplementary Fig. 1:** Hi-C heatmaps of diploid genomes of three *R. tenuis* accessions and *R. austrobrasiliensis*.

**Supplementary Fig. 2:** GenomeScope2 plots based on 31-mer of HiFi reads from all *R. tenuis* accessions as well as *R. austrobrasiliensis* and *R. breviuscula*.

**Supplementary Fig. 3:** Inferred tree by k-mer distance of autohexaploid *R. austrobrasiliensis* and the same fusion positions on both haplotypes of *R. tenuis*.

**Supplementary Fig. 4:** Probe distribution reveals absence of reciprocal translocations in *R. austrobrasiliensis*.

**Supplementary Fig. 5:** Meiotic chromosome behaviour of REC, PECP and JGV populations.

**Supplementary Fig. 6:** Distribution of 45S rDNA and REC translocation-specific probes (magenta - oligo-probe 1 and cyan - oligo-probe 2) during inverted meiosis in *R. tenuis* (PECP-48) visualised with FISH.

**Supplementary Fig. 7:** Pollen viability by Alexander Staining for each *R. tenuis* accession and *R. austrobrasiliensis* (percentage).

**Supplementary Fig. 8:** 3D rendering of an embryo sac from Toluidine blue-stained serial semithin physical sections stitched into a 3D z stack.

**Supplementary Fig. 9:** Pollen tube reaching the ovule.

**Supplementary Fig. 10:** Pollen tube reaching the mature embryo sac/female gametophyte to deliver the sperm cells.

**Supplementary Fig. 11:** Example of workflow for the genetic scan of meiotic genes.

**Supplementary Table 1.** Genome assembly statistics of all *R. tenuis* accessions and *R. austrobrasiliensis*.

| Species | Accession | Hap | Contig total length (bp) | Chr. total length (bp) | Contig number | Largest contig (bp) | Contig N50 (bp) | # gap | GC% |
|---|---|---|---|---|---|---|---|---|---|
| *R. tenuis* | REC | h1 | 395,484,736 | 348,158,620 | 1,663 | 31,615,665 | 19,340,799 | 28 | 35.68 |
| | | h2 | 373,542,889 | 354,512,391 | 616 | 71,890,041 | 23,642,622 | 24 | 35.48 |
| | JGV-16 | h1 | 512,049,716 | 377,634,235 | 1,408 | 112,179,635 | 61,459,675 | 7 | 39.41 |
| | | h2 | 380,901,205 | 349,658,151 | 145 | 66,517,139 | 36,225,668 | 18 | 35.18 |
| | JGV-17 | h1 | 573,037,334 | 377,385,800 | 2,032 | 179,483,847 | 46,422,628 | 11 | 41.90 |
| | | h2 | 440,003,022 | 348,979,334 | 447 | 61,337,029 | 35,842,360 | 9 | 38.34 |
| | JGV-89 | h1 | 502,650,004 | 377,247,412 | 2,119 | 65,727,966 | 54,108,842 | 18 | 40.04 |
| | | h2 | 368,703,004 | 349,182,916 | 240 | 51,407,378 | 36,217,777 | 28 | 36.36 |
| | PECP-47 | h1 | 420,138,026 | 381,933,207 | 945 | 72,058,618 | 37,276,100 | 13 | 36.62 |
| | | h2 | 359,408,259 | 349,003,858 | 208 | 84,674,538 | 38,765,659 | 13 | 35.32 |
| | PECP-48 | h1 | 550,339,684 | 382,648,587 | 3,562 | 66,123,649 | 7,984,712 | 46 | 38.25 |
| | | h2 | 387,787,195 | 350,935,085 | 585 | 33,381,499 | 12,378,200 | 63 | 36.21 |
| | PECP-35-6 | h1 | 426,271,194 | 382,455,145 | 871 | 141,969,674 | 75,026,608 | 12 | 36.47 |
| | | h2 | 360,109,026 | 350,070,891 | 158 | 55,915,206 | 38,097,382 | 15 | 35.48 |
| | PECP-36 | h1 | 394,251,559 | 382,926,756 | 304 | 72,005,413 | 26,891,453 | 27 | 35.33 |
| | | h2 | 378,334,318 | 348,977,203 | 963 | 60,647,232 | 32,395,499 | 20 | 35.33 |
| | PECP-36-7 | h1 | 414,955,118 | 383,444,050 | 651 | 111,369,649 | 50,763,498 | 12 | 35.78 |
| | | h2 | 379,302,196 | 349,631,601 | 605 | 66,243,789 | 38,781,351 | 15 | 35.34 |
| *R. austrobrasiliensis* | | | 2,168,431,183 | 2,117,633,834 | 1,136 | 42,774,656 | 10,796,733 | 212 | 35.55 |

**Supplementary Table 2: List of meiotic genes searched in *R. tenuis* genotypes, their location and comparison with *R. breviuscula* (Rb) and *R. austrobrasiliensis* (Ra).** The table reports several meiotic genes involved in recombination and 1) whether they were found as intact copies; 2) how many copies were found; 3) the species and haplotype they were found in; 4) a comparison with *R. breviuscula* and *R. austrobrasiliensis*; 5) references.

| Gene | REC hap1 | REC hap2 | JGV hap1 | JGV hap2 | PECP hap1 | PECP hap2 | Rb hap1 | Rb hap2 | Ra | RNA (REC) | Reference |
|---|---|---|---|---|---|---|---|---|---|---|---|
| SYN1/REC8 | 2 | 2 | 2 | 2 | 2 | 2 | 2 | 2 | 12 | + | 10.1371/journal.pgen.1010304 |
| SYN3 | 1 | 1 | 1 | 1 | 1 | 1 | 1 | 1 | 6 | + | 10.1007/s00412-009-0220-x |
| Top3alpha | 1 | 1 | 1 | 1 | 1 | 1 | 1 | 1 | 6 | + | 10.1371/journal.pgen.1000285 |
| HEIP1 | 1 | 1 | 1 | 1 | 1 | 1 | 1 | 1 | 6 | + | 10.1073/pnas.2221746120 |
| PMS1 | 1 | 1 | 1 | 1 | 1 | 1 | 1 | 1 | 6 | + | 10.1093/nar/gkaf187 |
| PHS1 | 1 | 1 | 1 | 1 | 1 | 1 | 1 | 1 | 6 | + | 10.1155/2012/514398 |
| PCH2 | 1 | 1 | 1 | 1 | 1 | 1 | 1 | 1 | 6 | + | 10.1093/nar/gkac1160 |
| SCEP2 | 1 | 1 | 1 | 1 | 1 | 1 | 1 | 1 | 6 | + | 10.1038/s41477-023-01558-y |
| SPO22/ZIP4 | 1 | 1 | 1 | 1 | 1 | 1 | 1 | 1 | 6 | + | 10.1371/journal.pgen.0030083 |
| SPO11/1 | 1 | 1 | 1 | 1 | 1 | 1 | 1 | 1 | 6 | + | 10.1105/tpc.107.054817 |
| SPO11/2 | 1 | 1 | 1 | 1 | 1 | 1 | 1 | 1 | 6 | + | 10.1105/tpc.107.054817 |
| SPO11/3 | 1 | 1 | 1 | 1 | 1 | 1 | 1 | 1 | 6 | + | 10.1111/pbi.13189 |
| MLH3 | 1 | 1 | 1 | 1 | 1 | 1 | 1 | 1 | 6 | + | 10.1038/sj.emboj.7600992 |

| | | | | | | | | | | |
|---|---|---|---|---|---|---|---|---|---|---|
| **MRE11** | **1** | **1** | **1** | **1** | **1** | **1** | **1** | **1** | **6** | **+** | **10.1007/s00412-007-0147-z** |
| **NBS1** | **1** | **1** | **1** | **1** | **1** | **1** | **1** | **1** | **6** | **+** | **10.1111/j.1365-313X.2007.03220.x** |
| **PRD2** | **1** | **1** | **1** | **1** | **1** | **1** | **1** | **1** | **6** | **+** | **10.1371/journal.pgen.1000654** |
| **PRD3** | **1** | **1** | **1** | **1** | **1** | **1** | **1** | **1** | **6** | **+** | **10.1371/journal.pgen.1010298** |
| **FANCM** | **1** | **1** | **1** | **1** | **1** | **1** | **1** | **1** | **6** | **+** | **10.1093/nar/gkac1244** |
| **MUS81** | **1** | **1** | **1** | **1** | **1** | **1** | **1** | **1** | **6** | **+** | **10.1073/pnas.2107543118** |
| **RAD50** | **1** | **1** | **1** | **1** | **1** | **1** | **1** | **1** | **7** | **+** | **10.1073/pnas.0906273106** |
| **RMI1** | **1** | **1** | **1** | **1** | **1** | **1** | **1** | **1** | **6** | **+** | **10.1093/nar/gkt730** |
| **RPA1a** | **1** | **1** | **1** | **1** | **1** | **1** | **1** | **1** | **6** | **+** | **10.3389/fpls.2016.00033** |
| **SEND1** | **1** | **1** | **1** | **1** | **1** | **1** | **1** | **1** | **7** | **+** | **10.1104/pp.114.237834** |
| **DFO** | **1** | **1** | **1** | **1** | **1** | **1** | **1** | **1** | **6** | **+** | **10.1111/j.1365-313X.2012.05075.x** |
| **ASY1** | **2** | **2** | **2** | **2** | **2** | **2** | **1** | **1** | **6** | **+** | **10.1093/pnasnexus/pgac302** |
| **ASY3** | **1** | **1** | **1** | **1** | **1** | **1** | **1** | **1** | **6** | **+** | **10.1104/pp.17.01725** |
| **ASY4** | **2** | **1** | **2** | **1** | **2** | **1** | **2** | **2** | **6** | **+** | **10.1104/pp.17.01725** |
| **DMC1** | **1** | **1** | **1** | **1** | **1** | **1** | **1** | **1** | **6** | **+** | **10.3389/fpls.2020.00839** |
| **HEI10** | **2** | **2** | **2** | **2** | **2** | **2** | **3** | **3** | **12** | **+** | **10.7554/eLife.79408** |
| **MER3** | **1** | **1** | **1** | **1** | **1** | **1** | **1** | **1** | **6** | **+** | **10.1371/journal.pone.0150482** |

| | | | | | | | | | | |
|---|---|---|---|---|---|---|---|---|---|---|
| **MLH1** | **1** | **1** | **1** | **1** | **1** | **1** | **1** | **1** | **6** | + | **10.1371/journal.pgen.1011197** |
| **MSH4** | **1** | **1** | **1** | **1** | **1** | **1** | **1** | **1** | **5** | + | **10.1371/journal.pgen.1003922** |
| **MSH5** | **2** | **1** | **2** | **1** | **2** | **2** | **1** | **1** | **9** | + | **10.1093/mp/sss145** |
| **SHOC1** | **1** | **1** | **1** | **1** | **1** | **1** | **1** | **1** | **6** | + | **10.1242/jcs.088229** |
| **ZYP1** | **1** | **3** | **1** | **1** | **1** | **1** | **1** | **1** | **6** | + | **10.1073/pnas.2021671118** |
| **RAD51** | **1** | **1** | **1** | **1** | **1** | **1** | **1** | **1** | **6** | + | **10.26508/lsa.202402701** |
| **MTOPVIb** | **1** | **1** | **1** | **1** | **1** | **1** | **1** | **1** | **6** | + | **10.1093/nar/gkae181** |
| **PRD1** | **2** | **2** | **2** | **2** | **2** | **2** | **2** | **2** | **12** | + | **10.1038/s41598-017-10270-9** |
| **HAP2** | **1** | **1** | **1** | **1** | **1** | **1** | **1** | **1** | **6** | + | **https://doi.org/10.1038/s41467-023-42789-z** |
| **NUF2** | **2** | **2** | **2** | **2** | **2** | **2** | **1** | **1** | **12** | + | **10.1111/tpj.15347** |
| **SCEP3** | **1** | **1** | **1** | **1** | **1** | **1** | **1** | **1** | **6** | + | **10.1038/s41477-025-02030-9** |
| **COMET** | **1** | **1** | **1** | **1** | **1** | **1** | **1** | **1** | **6** | + | **10.1016/j.cub.2020.07.089** |
| **SCEP1** | **1** | **1** | **1** | **1** | **1** | **1** | **1** | **1** | **6** | + | **10.1038/s41477-023-01558-y** |
| **BRCA2** | **1** | **1** | **1** | **1** | **1** | **1** | **1** | **1** | **6** | + | **PubMed: 38213474** |
| **PTD** | **1** | **1** | **1** | **1** | **1** | **1** | **1** | **1** | **6** | + | **10.1016/j.jgg.2014.02.001** |
| **RECQ4** | **1** | **1** | **1** | **1** | **1** | **1** | **1** | **1** | **6** | + | **10.1073/pnas.1713071115** |
| **MCC1** | **1** | **1** | **1** | **1** | **1** | **1** | **1** | **1** | **6** | + | **10.1111/j.1365-313X.2010.04191.x** |

**Supplementary Table 3.** Cell number of snRNA- and snATAC-seq of different *R. tenuis* accessions.

| **Source** | **Accession** | **Library** | **snRNA-seq** | **snATAC-seq** |
|---|---|---|---|---|
| Pollen | REC | 4984.A | 1,816 | x |
| | | 6067.C | x | 3,296 |
| | | 6067.D | x | 2,131 |
| | JGV-16 | 6451.C | 976 | x |
| | JGV-17 | 6451.B | 844 | x |
| | PECP-47 | 6385.B | 1,029 | x |
| | PECP-48 | 6451.A | 905 | x |
| Seed Embryo | REC | 7248.B | x | 327 |
| | PECP-48 | 7470.B | x | 185 |
| Seed Endosperm | REC | 7470.A | x | 3,365 |
| | PECP-48 | 7470.C | x | 76 |

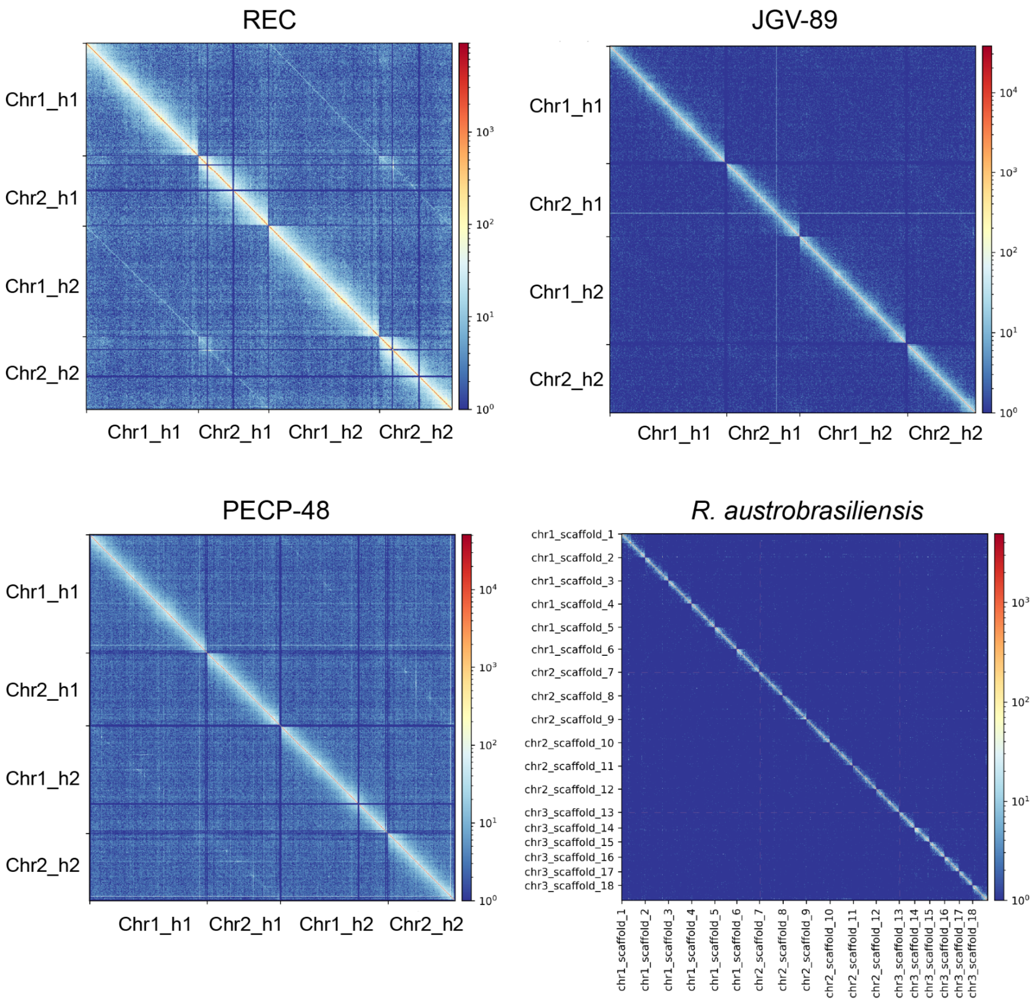


**Supplementary Fig. 1: Hi-C heatmaps of diploid genomes of three *R. tenuis* accessions and *R. austrobrasiliensis*.** The Hi-C maps of three *R. tenuis* accessions used 1Mb bin sizes. *R. austrobrasiliensis* used 2.5Mb resolution for plotting. The color bar on the right side of each Hi-C map indicates the Hi-C signal density.

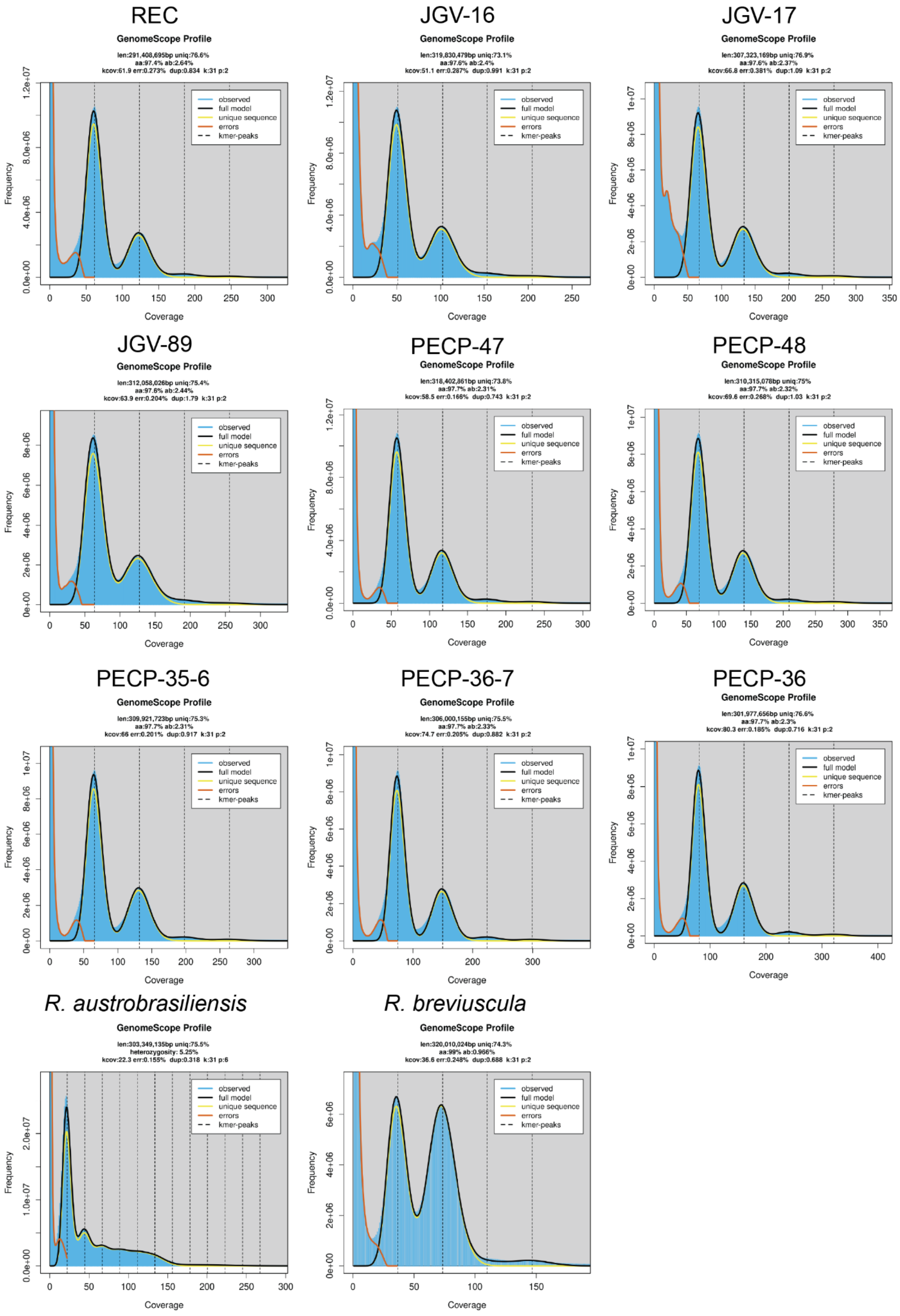

**Supplementary Fig. 2: GenomeScope2 plots based on 31-mer of HiFi reads from all *R. tenuis* accessions as well as *R. austrobrasiliensis* and *R. breviuscula*.**

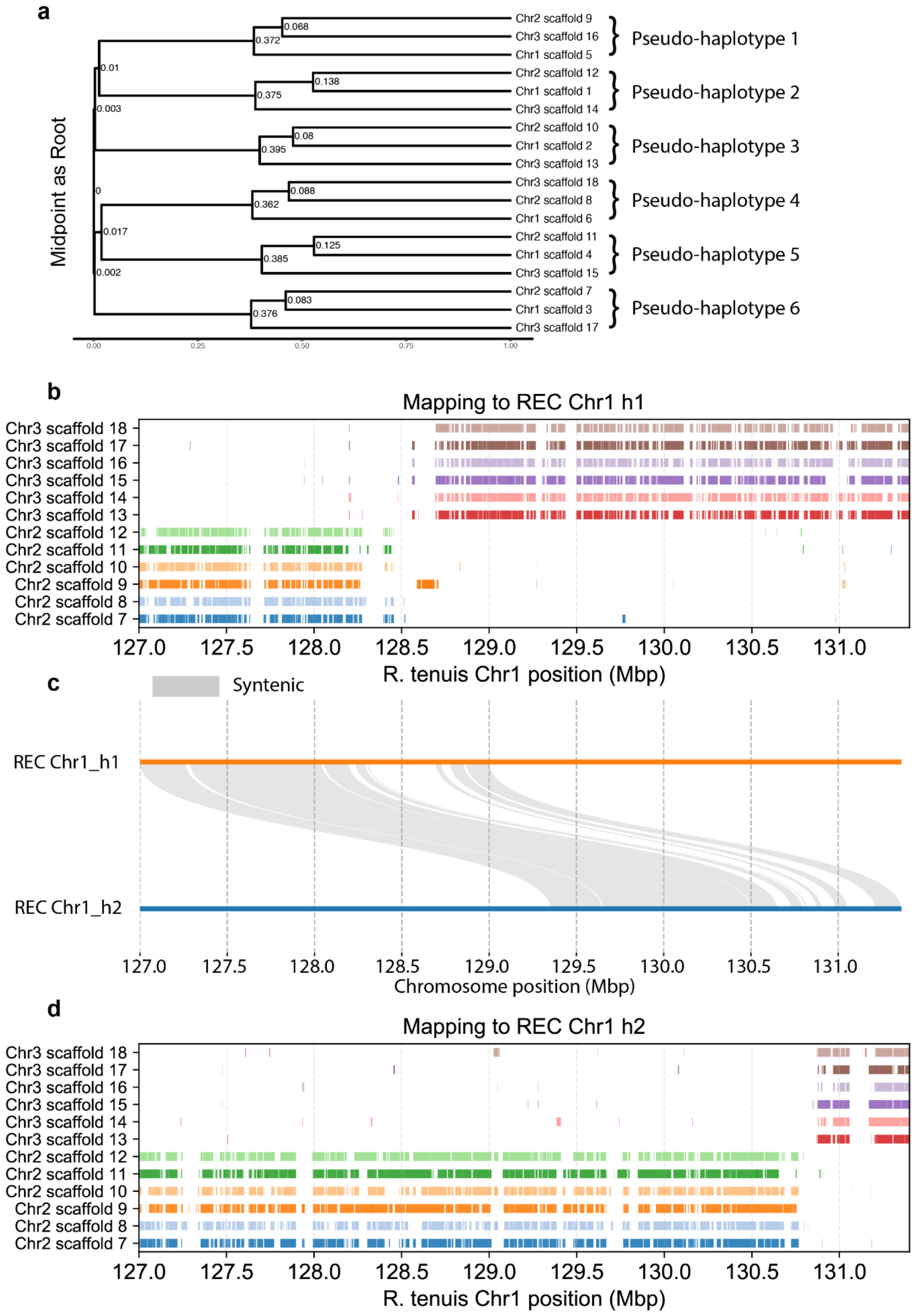


**Supplementary Fig. 3: Inferred tree by *K*-mer distance of autohexaploid *R. austrobrasiliensis* and the same fusion positions on both haplotypes of *R. tenuis*.** (**a**) Phylogenetic tree based on the k-mer distance matrix of all chromosome-level scaffolds of *R. austrobrasiliensis*. Labeled values are chronos (R function) scaled branch lengths assuming root edge has brach length 0 and sum of brach lengths from root to tip is 1. (**b**) Mapped fragments of *R. austrobrasiliensis* to *R. tenuis* (REC) haplotype 1 and (**d**) haplotype 2. (**c**) The SyRI collinearity of two haplotypes (REC) at the fusion positions.

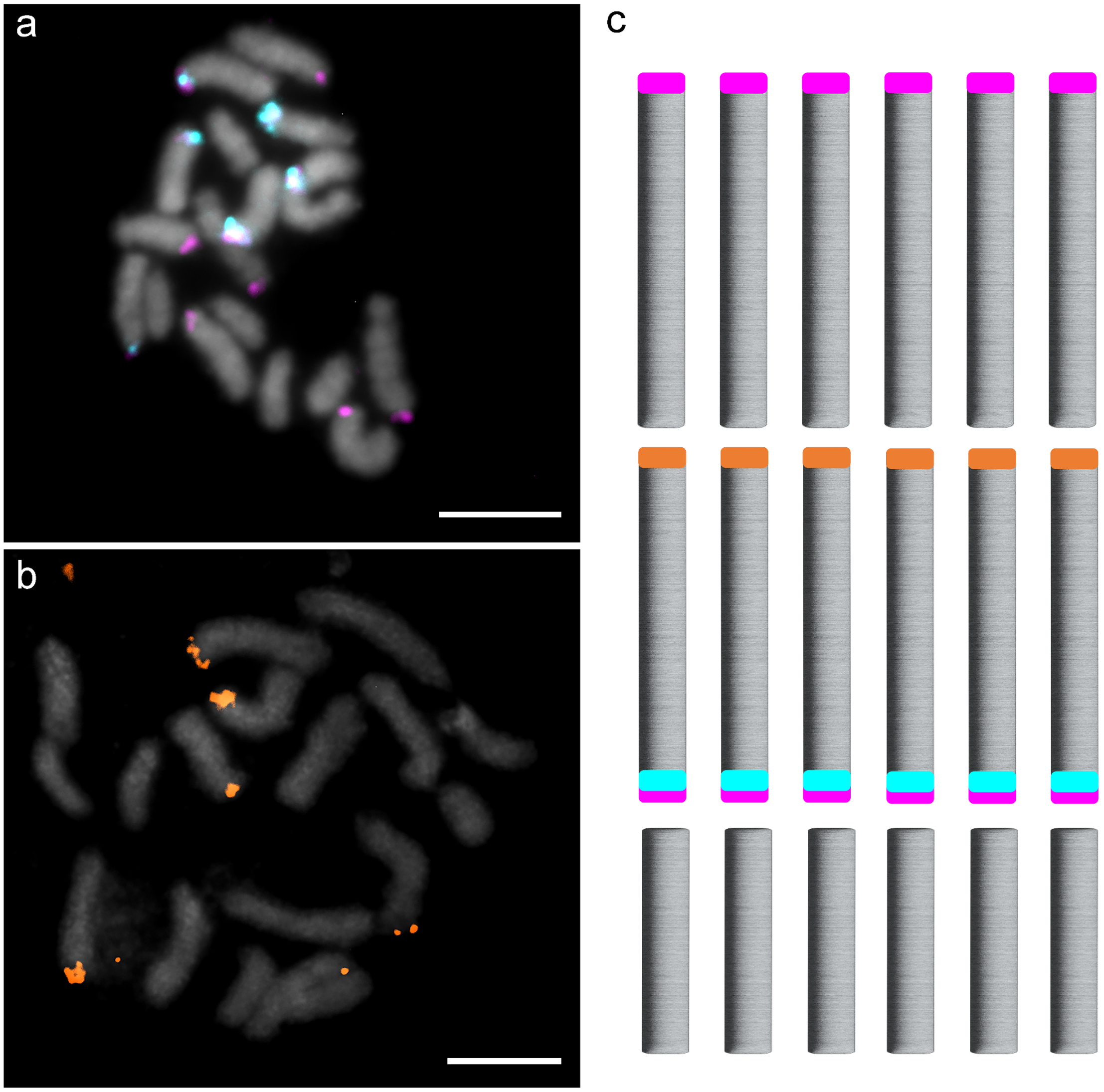


**Supplementary Fig. 4: Probe distribution reveals absence of reciprocal translocations in *R. austrobrasiliensis*.** FISH using translocation-specific probe sets (magenta, oligo-probe 1; cyan, oligo-probe 2) (**a**) and a 45S rDNA probe (orange) (**b**) on mitotic chromosomes of *R. austrobrasiliensis*. (**c**) Idiogram of *R. austrobrasiliensis* showing the distribution of probe signals from (**a**) and (**b**). The results indicate the absence of the reciprocal translocations found in *R. tenuis* genotypes. Scale bars correspond to 5 µm.

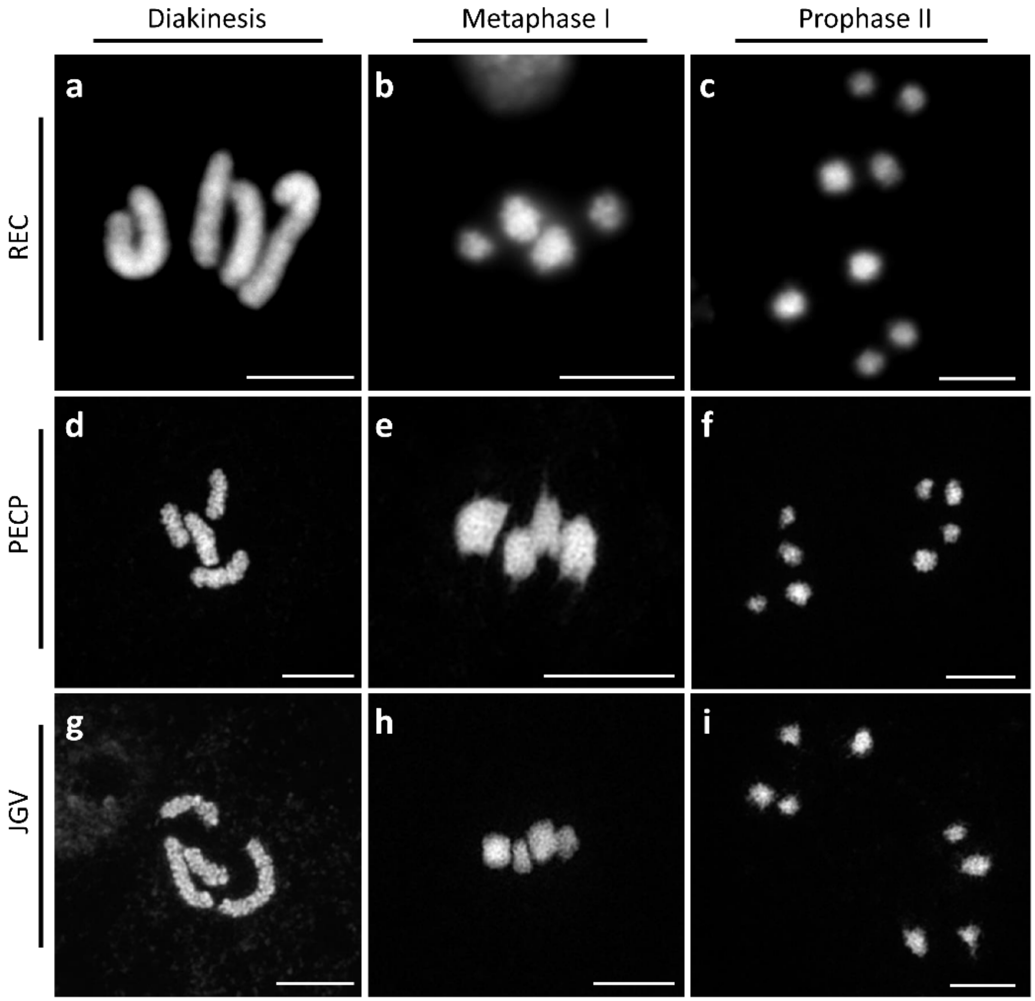


**Supplementary Fig. 5: Meiotic chromosome behaviour of REC, PECP and JGV populations**. At diakinesis, only 4 univalents are present and no bivalents are visible (**a**, n=10; **d**, n=21; **g**, n=40). Univalents align at metaphase I (**b**, n=15; **e**, n=16; **h**, n=11), followed by the segregation of sister chromatids that re-align at prophase II (**c**, n=22; **f**, n=26; **i**, n=12). Scale bars correspond to 5 µm.

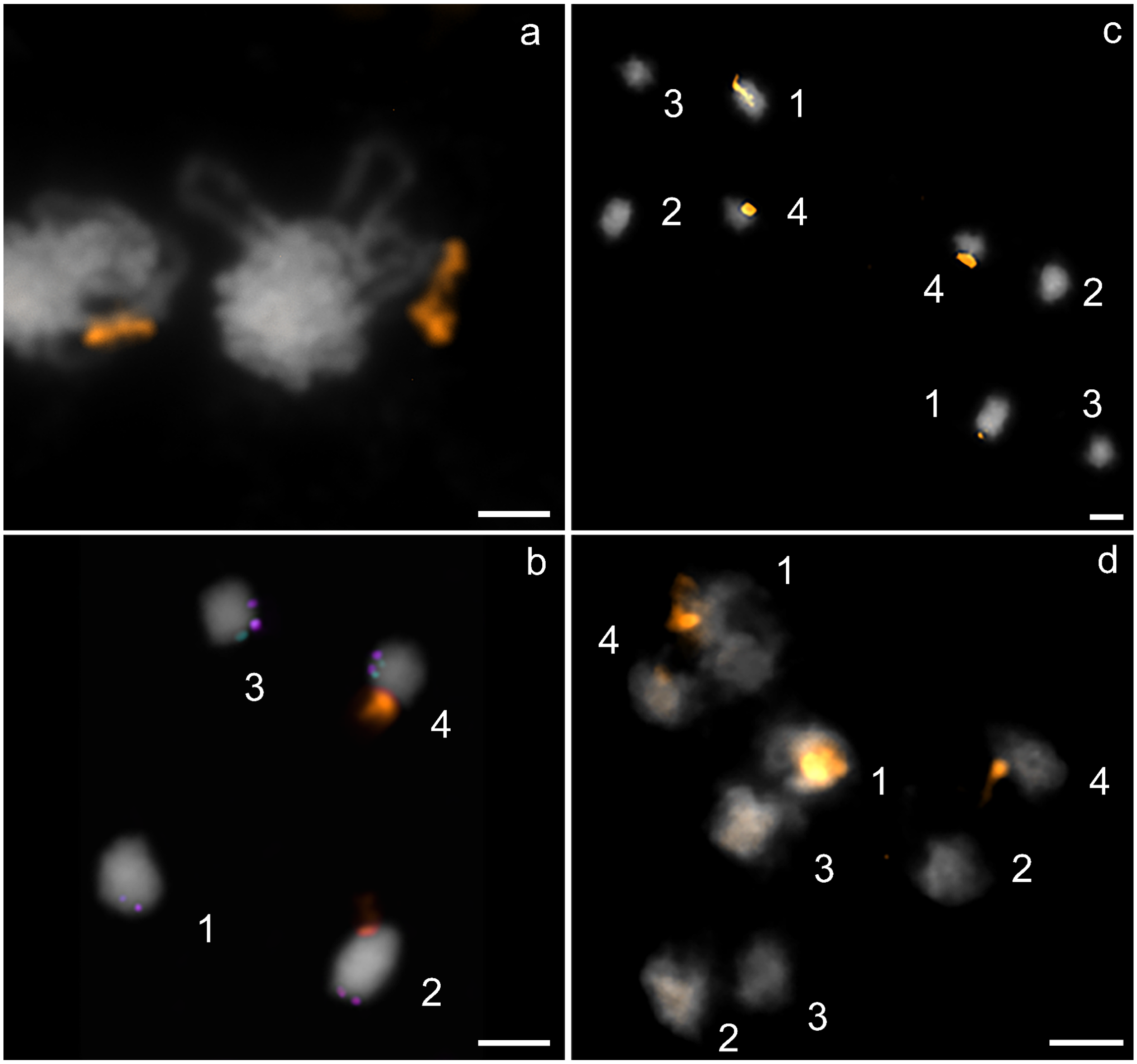


**Supplementary Fig. 6: Distribution of 45S rDNA and REC translocation-specific probes (magenta - oligo-probe 1 and cyan - oligo-probe 2) during inverted meiosis in *R. tenuis* (PECP-48) visualised with FISH.** (**a**) Zygotene, (**b**) Diakinesis, (**c**) Prophase II, (**d**) Anaphase II/Telophase II. 1 = Chr1_h1, 2 = Chr1_h2, 3 = Chr2_h1 and 4 = Chr2_h2. Note that the particular combination of 2 and 3 chromosomes in **d** does not harbour a NOR. Scale bars correspond to 2 μm.

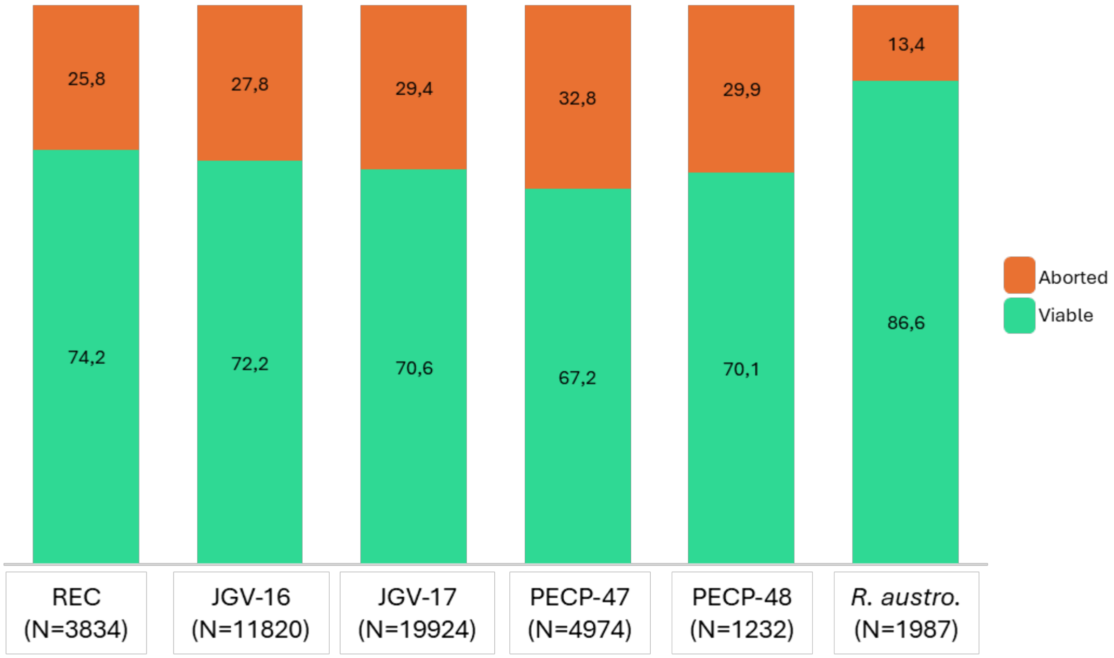


**Supplementary Fig. 7: Pollen viability by Alexander Staining for each *R. tenuis* accession and *R. austrobrasiliensis* (percentage).** Green indicates viable pollen, orange indicates aborted pollen. Total count (N) indicated in each accession label.

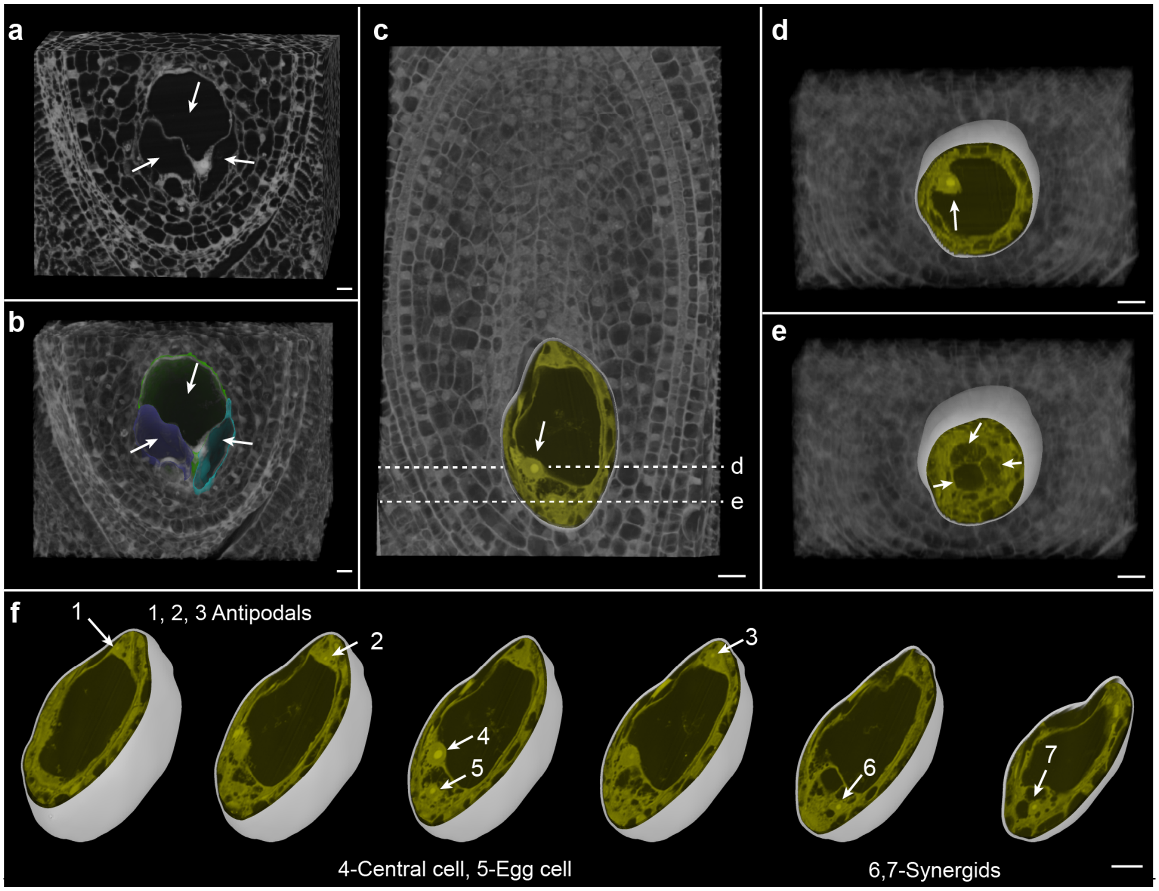


**Supplementary Fig. 8: 3D rendering of embryo sacs from Toluidine blue-stained serial semithin physical sections stitched into a 3D z stack.** (**a**–**b**) A case showing multiple embryo sacs in *R. tenuis*. Arrows point to the multiple embryo sacs (**b**) Embryo sac in (**a**) is visualized as a segmented object, with each one assigned a different color for distinction. (**c**) Longitudinal section view of a mature 7-nuclei embryo sac depicting the large central cell nucleus (arrow). Dotted lines represent the orientation of transverse sections shown in (**d**–**e**). (**d**) 3D cropped view of the sample along the transverse section showing the central cell nucleus (arrow). (**e**) 3D cropped view of the sample along the transverse section showing the egg apparatus containing the egg cell and the two adjacent synergids (arrow). (**f**) Series of cropped 3D views of the extracted embryo sac structure, showing the seven nuclei in different 2D planes through the mature embryo sac. (**c**–**f**) 3D segmentation of the embryo sac is shown as a white object, with the Toluidine blue-stained section overlaid in greyscale and the embryo sac region false-colored yellow. 3D rendering was performed using MorphoGraphX. Scale bar 10µm.

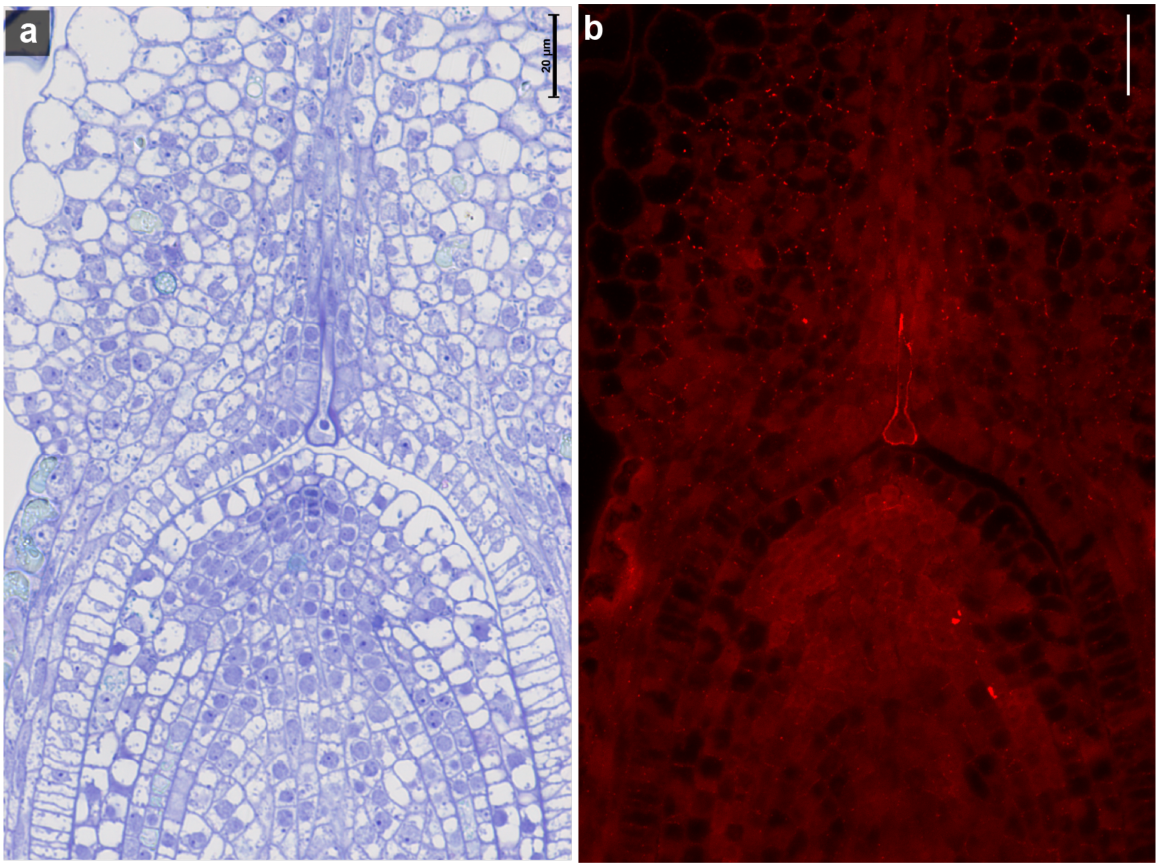


**Supplementary Fig. 9: Pollen tube reaching the ovule.** (**a**) Toluidine blue-stained semithin section depicting a pollen tube reaching the distal part of the ovule after growing down the style of the ovary. (**b**) Indirect immunofluorescent labeling of ß-1,3-glucan epitopes of the semithin section shown in (**a**) highlights the callose pollen tube cell wall in the surrounding ovary tissue. Note the punctate labeling of plasmodesmata decorating the cell walls in the beak-like cap tissue. Autofluorescence is caused by glutaraldehyde in the primary fixative and the previous Toluidine blue staining. Scale bar 20µm.

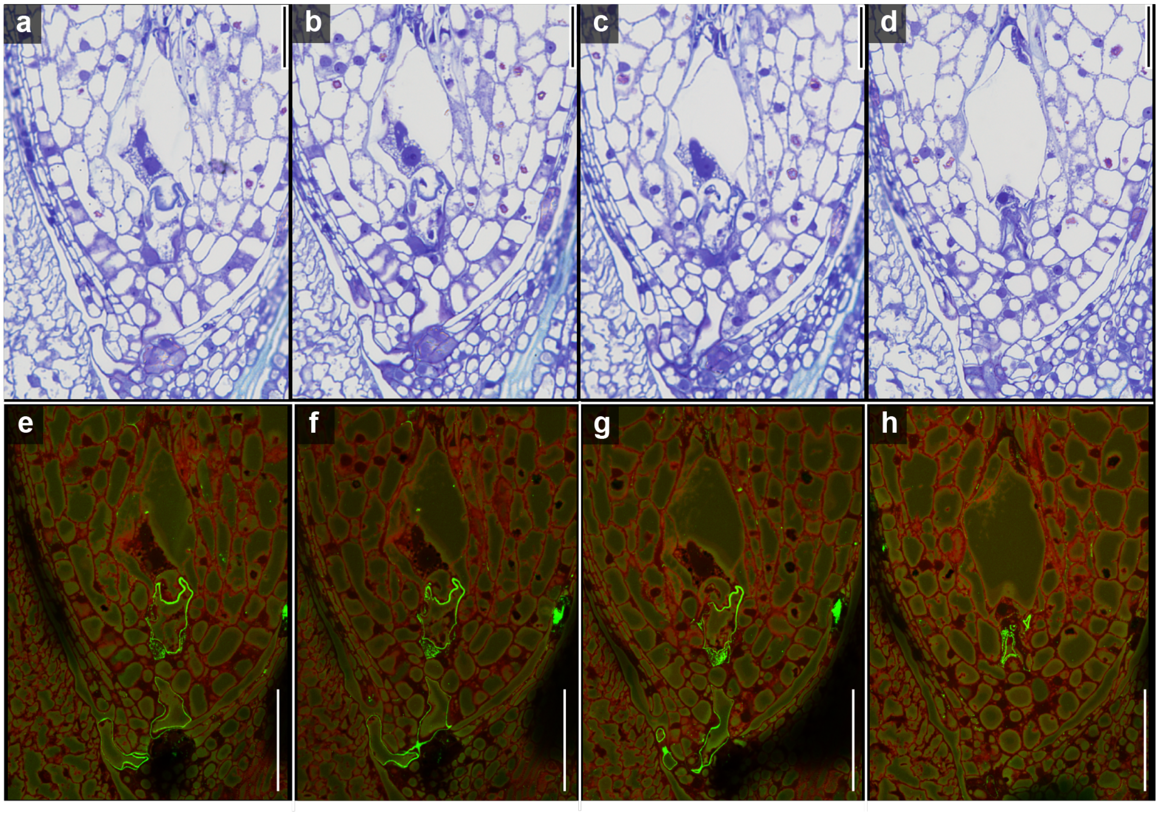


**Supplementary Fig. 10: Pollen tube reaching the mature embryo sac/female gametophyte to deliver the sperm cells.** (**a**–**d**) Consecutive toluidine blue-stained semithin sections depicting a pollen tube that has grown through the micropyle formed by the inner integument and reached the mature embryo sac. The globular nucleus of the central cell and - in direct contact - one of the two lens-shaped sperm nuclei can be seen. (**b**–**h**) Indirect immunofluorescent labeling of ß-1,3-glucan epitopes of the semithin sections shown in **a**–**g** highlights the callose pollen tube cell wall in the surrounding ovary tissue. Autofluorescence in both channels is caused by glutaraldehyde in the primary fixative and previous Toluidine blue staining. Scale bar 50µm.

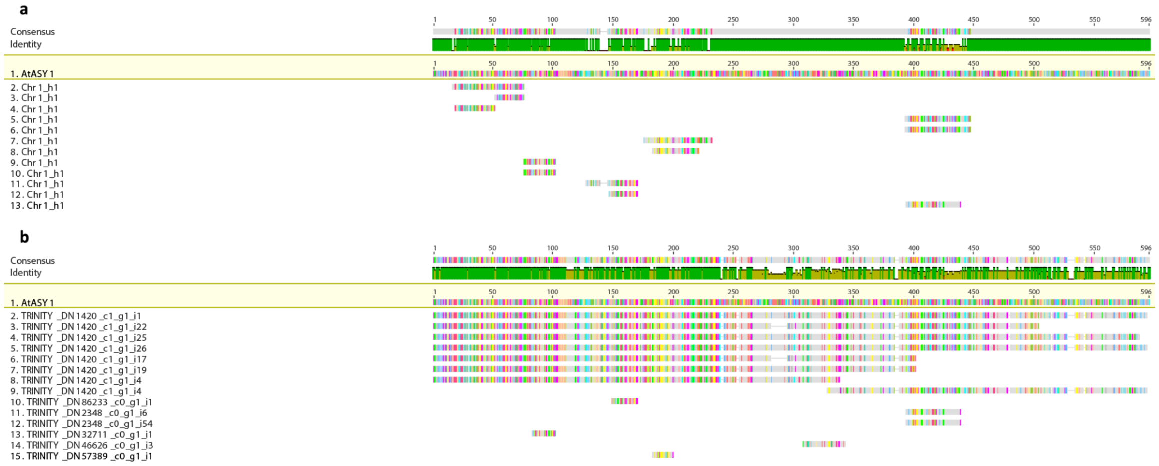


**Supplementary Fig. 11: Example of workflow for the genetic scan of meiotic genes.** (**a**) ASY1 shows two overlapping hits for each query region, suggesting a duplication. (**b**) ASY1 can be found in our anther-specific transcriptome. Manual curation revealed a tandem duplication (156.939.594 -> 156.948.842 and 184.686.149 -> 184.695.397) on chromosome 1.
